# Transfer learning between preclinical models and human tumors identifies conserved NK cell activation signature in anti-CTLA-4 responsive tumors

**DOI:** 10.1101/2020.05.31.125625

**Authors:** Emily F. Davis-Marcisak, Allison A. Fitzgerald, Michael D. Kessler, Ludmila Danilova, Elizabeth M. Jaffee, Neeha Zaidi, Louis M. Weiner, Elana J. Fertig

**Affiliations:** McKusick-Nathans Institute of the Department of Genetic Medicine, Johns Hopkins School of Medicine, Baltimore, MD, USA; Department of Oncology, Sidney Kimmel Comprehensive Cancer Center, Johns Hopkins School of Medicine, Baltimore, MD, USA; Department of Oncology, Georgetown Lombardi Comprehensive Cancer Center, Georgetown University Medical Center, Washington, DC, USA; Department of Applied Mathematics and Statistics, Johns Hopkins University Whiting School of Engineering, Baltimore, MD, USA; Department of Biomedical Engineering, Johns Hopkins University School of Medicine, Baltimore, MD, USA

## Abstract

**Background:** Tumor response to therapy is affected by both the cell types and the cell states present in the tumor microenvironment. This is true for many cancer treatments, including notably immune checkpoint inhibitors (ICIs). While it is well-established that ICIs promote T cell activation, their broader impact on other intratumoral immune cells is unclear; this information is needed to identify new mechanisms of action and improve ICI efficacy. Many preclinical studies have begun to use single cell analysis to delineate therapeutic responses in individual immune cell types within tumors. One major limitation to this approach is that therapeutic mechanisms identified in preclinical models have failed to fully translate to human disease, restraining efforts to improve ICI efficacy in bench to bedside research.

**Method:** We previously developed a computational transfer learning approach to identify shared biology between independent high-throughput single-cell RNA sequencing (scRNA-seq) datasets. In the present study, we test this framework’s ability to identify conserved and clinically relevant transcriptional changes in complex tumor scRNA-seq data and further expand its application beyond comparison of scRNA-seq datasets into comparison of scRNA-seq datasets with additional data types such as bulk RNA-seq and mass cytometry.

**Results:** We found a conserved signature of NK cell activation in anti-CTLA-4 responsive mice and human tumors. In human melanoma, we found that the NK cell activation signature correlates with longer overall survival and is predictive of anti-CTLA-4 (ipilimumab) response. Additional molecular approaches to confirm the computational findings demonstrated that human NK cells express CTLA-4 and bind anti-CTLA-4 independent of the antibody binding receptor (FcR), and that similar to T cells, CTLA-4 expression by NK cells is modified by cytokine-mediated and target cell-mediated NK cell activation.

**Conclusions:** These data demonstrate the ability of our transfer learning approach to identify cell state transitions conserved in preclinical models and human tumors. This approach can be adapted to explore many immuno-oncology questions, enhancing bench to bedside research and enabling better understanding and treatment of disease.

**Graphical Abstract:** 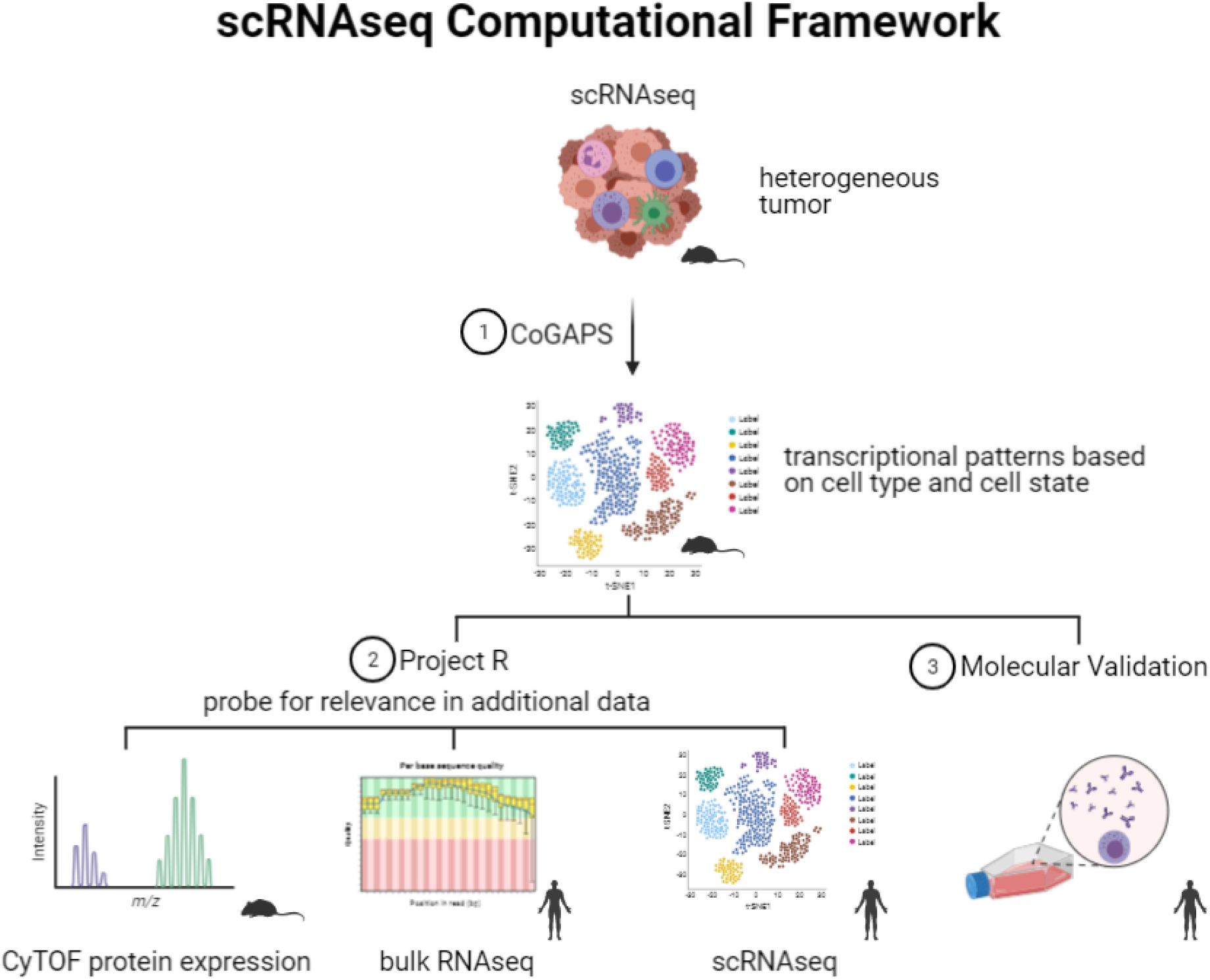

## 1. Introduction

Single-cell RNA-sequencing (scRNA-seq) provides an unprecedented opportunity to unravel the cellular complexity and diversity of immune cell populations in the tumor microenvironment[1].

When used in the context of immunotherapy, scRNA-seq can provide a more comprehensive understanding of the molecular and cellular pathways that drive therapeutic response and resistance. While studies often use preclinical mouse models as a convenient and useful tool for studying therapeutic response mechanisms, they are limited in their ability to infer biology relevant to therapeutic responses in humans. To improve the clinical efficacy of immunotherapies such as immune checkpoint inhibitors (ICIs), we need a deeper understanding of the fundamental mechanisms that underlie the anti-tumor activity of ICIs in humans.

Many aspects of the immune system are conserved between mice and humans, but there are significant species-specific differences[2]. These differences may contribute to the frequent failure of therapies that are effective in mouse models from showing similar efficacy in humans[3]. Discrepancies between ICI mechanisms observed in mice and humans may be further complicated by species-specific differences that mask detection of conserved alterations in responding immune cells. A deeper understanding of human and mouse immune responses to immunotherapy could generate new insights into properties that define therapeutic sensitivity.

Emerging scRNA-seq studies that have begun to characterize changes in gene expression after immunotherapy treatment[4–6] are ideally suited to begin learning these mechanisms. In order to accomplish this, computational tools that identify conserved cell state transitions across species are needed to compensate for species-specific immune system differences in transcriptional data. As scRNA-seq becomes increasingly popular in immuno-oncology, such tools will be essential to validate preclinical computational findings in terms of both robustness and clinical relevance.

Recently, we developed a computational framework that uses matrix factorization and transfer learning to integrate transcriptional datasets from different species[7]. This has led to the identification of both species-specific and conserved biological processes in the developing retina of mice and humans[8,9]. In the context of cancer, this framework has the potential to identify complex cellular alterations within the tumor microenvironment induced by therapy. In this study, we use this framework’s ability to identify conserved and clinically relevant transcriptional changes in scRNA-seq data of immune cells from ICI treated tumors. We further compare biological features across additional data types such as bulk RNA-seq and mass cytometry. We demonstrate the ability of our framework to identify shared tumor immune biology present across independent datasets derived from different tumor types, treatment groups, sequencing platforms, and species. We detect a robust signature of NK cell activation that is associated with positive clinical outcomes in response to anti-CTLA-4 and overall survival in treatment-naive tumors. We confirm the relevance of our computational findings by using molecular techniques to begin elucidating how NK cells are activated in response to anti-CTLA-4 treatment. These analyses yield novel insights into the role of NK cells in anti-CTLA-4 efficacy and provide computational tools that can be applied to other therapeutic datasets to enable translational cancer immunotherapy research.

## 2. Results

### CoGAPS identifies known molecular alterations in response to immunotherapy from scRNA-seq data

To detect transcriptional signatures (also called “patterns”) that represent biological features across intratumoral immune cells during immunotherapy response, we used our non-negative matrix factorization (NMF) technique, CoGAPS (Fig 1A)[10]. CoGAPS is an established approach to dissect transcriptional signatures that dictate cell type identity (i.e., NK vs. Treg) and cell state (i.e., activated vs. resting), aiding the evaluation of complex molecular alterations within the tumor immune microenvironment[11,12]. By combining CoGAPS with projectR, a transfer learning approach, we can then quickly query for shared features across independent datasets (Fig. 1A)[7,10].

**Figure 1.**
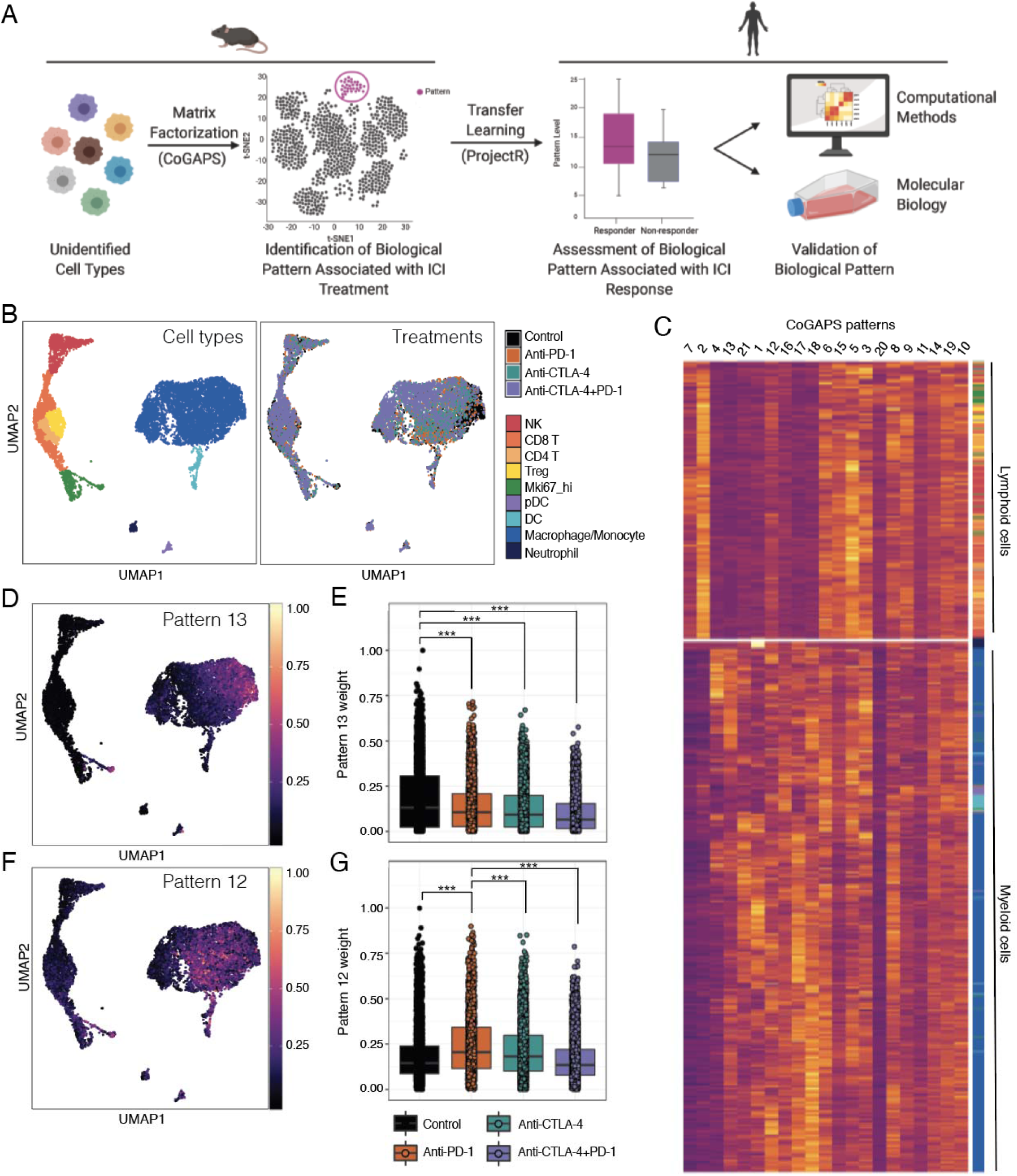
CoGAPS identifies gene signatures related to immune cell lineage and treatment response in mouse intratumoral immune cell scRNA-seq data. **A.** We apply CoGAPS, a non-negative matrix factorization algorithm, to publicly available scRNA-seq data of ICI-treated mouse tumors. Matrix factorization algorithms are unsupervised learning methods that can distinguish the molecular dynamics of therapeutic responses without prior knowledge of gene regulation or cell type classification. Using the transfer learning method projectR, the transcriptional signatures (or patterns) identified by CoGAPS are then projected into an independent dataset of human tumors treated with ICIs. These signatures can then be computationally assessed for relationships to clinical outcomes and molecularly validated in human cell lines. **B.** UMAP-dimension reduction of droplet-based scRNA-seq of intratumoral immune cells from ICI treated mouse sarcomas[4]. Samples are colored by annotated cell types (left) and by treatment (right). **C.** Hierarchical clustered heatmap of 21 CoGAPS patterns demonstrating segregation by immune cell lineage. Rows are individual cells, with column annotations designating cell type. Columns represent different CoGAPS patterns. **D.** UMAP-dimension reduction colored by CoGAPS pattern 13 weights illustrates a cell type specific signature within the macrophages/monocytes. **E.** Boxplot of pattern 13 weights in individual macrophage/monocyte cells, faceted by treatment group. Pattern 13 is associated with cells treated with control monoclonal antibody. **F.** UMAP-dimension reduction colored by CoGAPS pattern 12 weights illustrates a cell type specific signature within the macrophages/monocytes. **G.** Boxplot of pattern 12 weights in individual macrophage/monocyte cells, faceted by treatment group. Pattern 12 is associated with cells treated with anti-PD-1.

To identify transcriptional responses induced by ICIs in mouse tumors, we applied CoGAPS to a publicly available scRNA-seq dataset including more than 15,000 immune cells isolated from mouse sarcomas[4]. These tumors were treated with a control monoclonal antibody, anti-PD-1, anti-CTLA-4, or combination anti-PD-1 and anti-CTLA-4 antibodies (Fig. 1B). A critical challenge in matrix factorization algorithms such as CoGAPS is the selection of an appropriate dimensionality (i.e., number of patterns) to resolve biological features from the data[13]. Consistent with previous studies, running CoGAPS across multiple-dimensionalities revealed that different levels of biological complexity were captured at different dimensionalities[14]. For example, at low dimensionality (3 patterns) CoGAPS separated immune cells into myeloid and lymphoid lineages (Supplemental Fig. 1A). When dimensionality was increased to 21 patterns, the myeloid versus lymphoid lineage distinction was preserved and additional transcriptional signatures reflecting immune cell type and state were captured (Fig. 1C). To identify specific attributes captured by each pattern, we performed gene set analysis using the gene weights for each pattern as input. We used the hallmark gene sets from the Molecular Signatures Database (MSigB)[15] and the PanCancer Immune Profiling gene panel from Nanostring Technologies to assess enrichment of gene sets controlling well-defined biological processes. Gene set statistics for all patterns are provided in supplemental Table 1.

We found that several transcriptional signatures identified by CoGAPS were consistent with ICI-mediated changes previously described in the literature. For example, pattern 13 was enriched in macrophages/monocytes from progressing tumors treated with control monoclonal antibody (Fig. 1D and E) while pattern 12 was prevalent in macrophages/monocytes from tumors treated with anti-PD-1 (Fig. 1F and G). Macrophages are commonly divided into two subsets, pro-inflammatory anti-tumor M1 subtype and anti-inflammatory pro-tumor M2 subtype[16]. Consistent with this, pattern 13, which was enriched in control-treated tumors, reflected M2 macrophage polarization, which promotes tumor growth and metastasis. In contrast, pattern 12, which was enriched in anti-PD-1 treated tumors, reflected M1 macrophage polarization and interferon responses. This finding agrees with a recent study, which showed that anti-PD-1 treatment leads to a functional transition within the macrophage compartment towards an immunostimulatory M1 phenotype[17].

### CoGAPS analysis identifies a subset of activated NK cells in mouse tumors treated with anti-CTLA-4

In addition to the known transcriptional changes shown in Figure 1, CoGAPS also identified a transcriptional signature that reflected a subset of activated NK cells—pattern 7 (Fig. 2A and B). While tumors from each treatment group contained NK cells with elevated levels of pattern 7, there was a significant enrichment in NK cells from tumors that were treated with anti-CTLA-4 (Fig. 2C). To identify genes strongly associated with this pattern, we used the CoGAPS PatternMarker statistic[18]. PatternMarker analysis identified 3,195 genes associated with pattern 7. Gene set enrichment analysis of these genes revealed an upregulation of interferon-gamma and IL2-STAT5 gene sets, which are key pathways that govern cytotoxicity and maturation in NK cells (Supplemental Table 1)[19].

**Figure 2.**
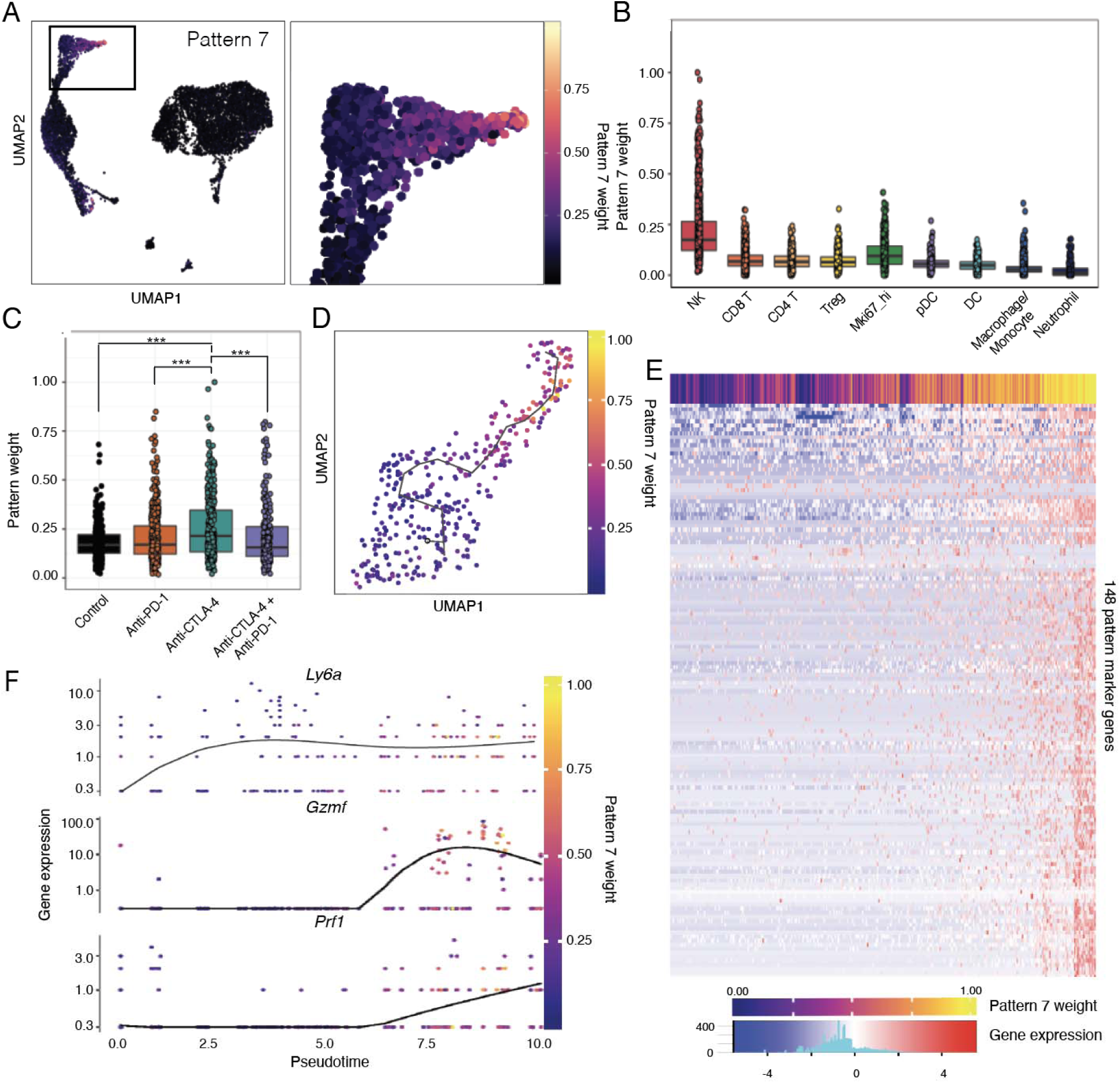
CoGAPS and pseudotime analysis reveals a dynamic state change in NK cells during ICI exposure in mouse scRNA-seq data. **A.** UMAP dimension reduction colored by CoGAPS pattern 7 weights across all cells (left) and magnified view (right) showing that pattern 7 marks a population of NK cells delineated in Fig. 1A. **B.** Boxplot of pattern 7 weights across each immune cell type. Cells with high pattern 7 weights are observed only in NK cells. **C.** Boxplot of pattern 7 weights in individual NK cells faceted by treatment group. Anti-CTLA-4 treated NK cells have increased pattern 7 weights compared to NK cells treated with other immunotherapies. **D.** Pseudotemporal trajectory of anti-CTLA-4 treated NK cells colored by CoGAPS pattern 7 weight suggesting that anti-CTLA-4 treatment results in NK cell activation. **E.** Heatmap of gene expression for 148 pattern markers that are differentially expressed across pseudotime. Columns are individual cells, and column annotation designates pattern 7 weight in each cell. Rows are differentially expressed pattern markers. **F.** Gene expression of selected NK cell activation genes differentially upregulated across pseudotime. Each dot represents a different cell and is colored by CoGAPS pattern 7 weight.

We hypothesized that pattern 7 was identifying NK cells undergoing a cell state change in response to ICI. To explore this possibility we performed pseudotime analysis on NK cells from tumors treated with anti-CTLA-4[20]. Pseudotime analysis enables a quantitative estimation of cellular progression through dynamic biological processes. The pseudotemporal ordering showed a sequential progression in cellular trajectory during anti-CTLA-4 treatment (Figure 2D). This pseudotemporal trajectory was highly correlated with the pattern 7 weight identified in each cell (0.71 spearman correlation). Notably, the trajectory revealed a single transition state in NK cells as a result of anti-CTLA-4 treatment, with individual cells having transcriptional profiles that reflect various points along the trajectory. Differential expression analysis across pseudotime identified 1,968 genes with significant changes (q value < 0.01) in gene expression during exposure to anti-CTLA-4 (Supplemental Table 2). We then looked for differentially expressed genes over pseudotime that were strongly associated with pattern 7 as determined by patternMarker analysis (Fig. 2E). The 148 differentially expressed patternMarker genes included markers of NK cell activation, including as perforin, granzymes, and Ly6a[21], that significantly increased in expression along the pseudotime trajectory as a result of anti-CTLA-4 treatment (Fig. 2F). These data support recent findings that NK cells within mouse tumors can be functionally modulated by ICI treatment[22,23].

In their original study, Gubin et al. used CyTOF, a mass spectrometry-based flow cytometry method to measure protein expression, in parallel with their scRNA-seq. By CyTOF they found that anti-CTLA-4 induced Granzyme B in a population of KLRG1+ NK cells. While these KLRG1+ NK cells resembled a population of NK cells detected by scRNA-seq, the relationship between anti-CTLA-4 and NK cell activation was unclear. We hypothesized that the KLRG1+ NK cells identified by CyTOF would contain the transcriptional NK cell activation signature we detected by scRNA-seq. To test this hypothesis, we used our transfer learning method, projectR [24], to assess the CyTOF data for the 21 patterns identified by CoGAPS from scRNA-seq. Just as with the scRNA-seq data, pattern 7 was highest in lymphocytes from anti-CTLA-4 treated tumors in the CyTOF data (Supplemental Fig. 1B). This demonstrates that: 1) CoGAPS is able to identify transcriptional changes in response to immunotherapy, which are preserved at the protein and mRNA level and across technological platforms and 2) CoGAPS identified an NK cell activation signature in the scRNA-seq data that was missed by the traditional scRNA-seq analysis methods used in the original study and 3) ProjectR is capable of identifying gene expression signatures present in scRNA-seq and CyTOF data.

### Preclinical NK cell activation signature is associated with overall in metastatic melanoma patients

To evaluate the clinical relevance of the NK cell activation signature (pattern 7) and the ability of ProjectR to identify conserved biological processes in mouse and human tumors, we projected bulk RNA-seq data from 9,553 untreated human tumors representing 32 cancer types onto the 21 mouse patterns[25]. This enabled a pan-cancer investigation of the relationship between the mouse tumor immune cell signatures identified by CoGAPS and clinical outcomes in human disease. We fit a multiple linear regression model to estimate the association between the projected weight of each pattern and overall survival. When including cancer type as a covariate in the model given its significant effect on survival, we found that the NK cell activation signature was the most significantly associated with overall survival, as compared to the other patterns (Fig. 3A, p < 6 x 10^−5^). Pattern 15, which was similarly associated with mouse NK cells, was also significantly associated with overall survival (Supplemental Fig. 2A, Fig. 2B, p < 5.9 x 10^−4^). When including age as a covariate in the linear model, the NK cell activation signature remains the most significantly associated with overall survival (Supplemental Fig. 2B, p < 1.6 x 10^−4^). Interestingly, the NK cell activation signature was the only pattern to show a significant negative correlation with age (Supplemental Fig. 2C, p < 6.7 x 10^−3^). Several studies have reported age-related alterations in NK cell function, including a decreased ability to proliferate and kill target cells in older individuals[26,27]. The NK cell activation signature appears to decrease as individuals age, which may have implications for cancer incidence in elderly individuals.

**Figure 3.**
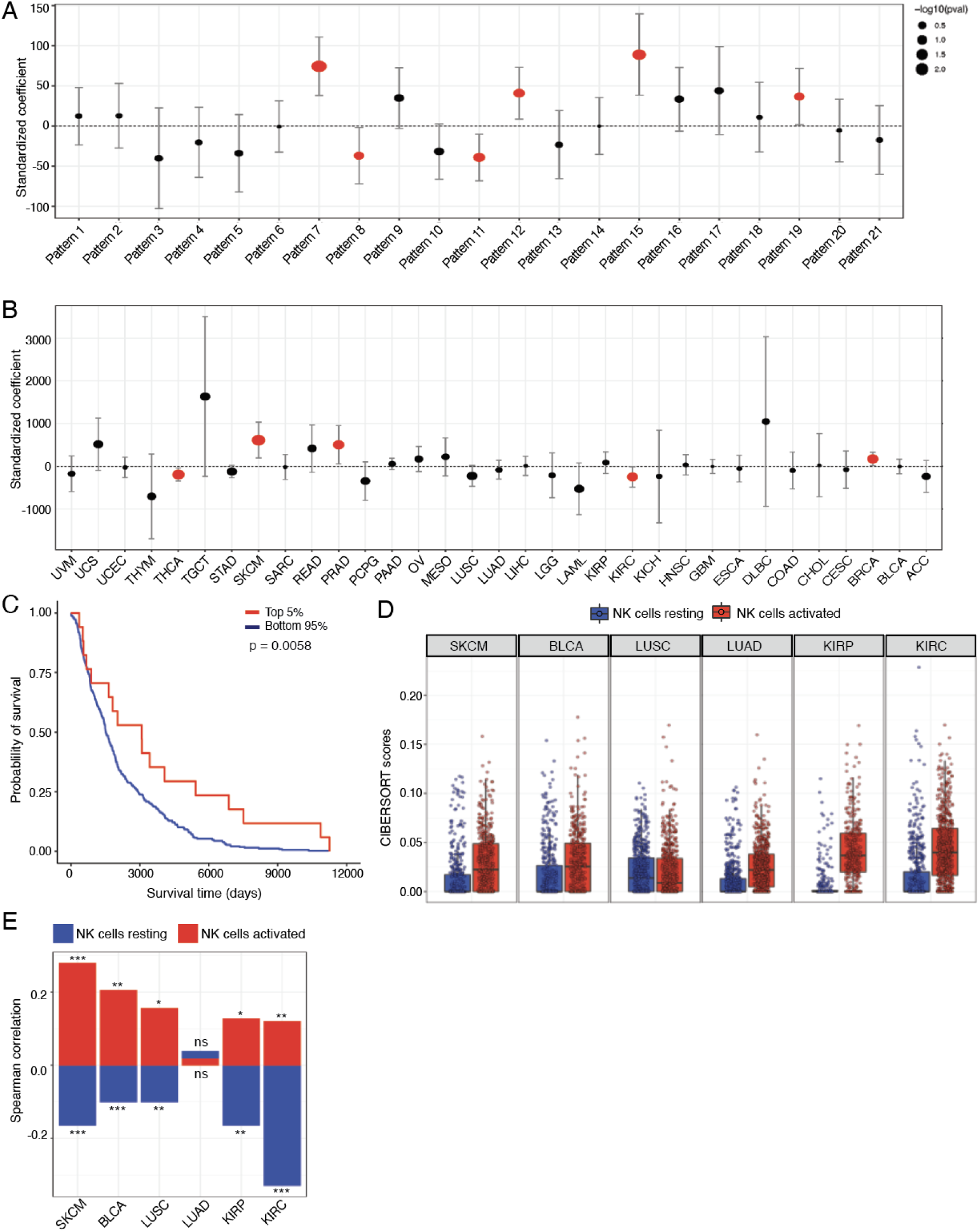
Preclinical NK activation signature is associated with overall survival in human melanoma. **A.** The output from a multiple linear regression model that predicts overall tumor survival from the CoGAPS transcriptional signatures, while also adjusting for cancer type as a covariate. Standardized coefficients (i.e. data was scaled and centered) representing the strength and direction of association for each pattern are shown on the x-axis, with error bars representing coefficient +− 1.96 * standard error, and point size scaled to the coefficient’s p-value. Patterns 7 and 15 are most strongly associated with longer overall survival, with pattern 7 being most significantly positively associated (p < 1.2 x 10^−4^). **B.** The output from a multiple linear regression model that predicts overall tumor survival from the CoGAPS transcriptional signatures, while also adjusting for patient age as a covariate. Pattern 7 is the most significantly positively associated with overall survival in SKCM (p < 5 x 10^−3^). **C.** Kaplan-Meier plot of overall survival for 368 metastatic melanoma patients with the top 5% and bottom 95% of pattern 7 scores. **D.** Boxplot of CIBERSORT scores estimating the abundance of resting and activated NK cells from TCGA RNA-seq data by tumor subtype in TCGA. **E.** Bar plot of Spearman correlation coefficients between CTLA-4 and CIBERSORT cell type score for immunogenic cancers. CTLA-4 expression is positively correlated with estimation of activated NK cells from TCGA RNA-seq data. Significant correlations for NK scores and CTLA-4 expression are indicated by asterisks where p-values < 0.05 = *, < 0.01 = **, and p-values < 0.001 = ***.

When fitting separate regression models by cancer type, we found that melanoma (SKCM) had the strongest and most significant association between the NK cell activation signature and longer overall survival (Fig. 3B, Supplemental Fig. 2D, p < 0.005). Notably, this association was driven entirely by the metastatic melanoma samples (Supplemental Fig. 2E, F), which is consistent with the role of NK cells controlling cancer progression and metastasis[28]. Melanoma patients with tumors that had elevated NK cell activation signature (top 5%) had significantly longer overall survival (Fig. 3C). Prostate cancer (PRAD) and breast cancer (BRCA) also showed a positive correlation between increased NK cell activation signature and longer overall survival (Fig. 3B). These results demonstrate we can computationally identify transcriptional signatures relevant to clinical outcomes from preclinical mouse datasets and confirm that NK cell activation is associated with overall survival in metastatic melanoma[29].

### CTLA-4 expression is positively correlated with the infiltration of active NK cells in immunogenic human tumors

Given that the NK cell activation signature was enriched in anti-CTLA-4 treated mouse tumors, we hypothesized that there may be a correlation between CTLA-4 expression and intratumoral NK cell content. To explore this hypothesis, we used bulk-RNA-seq data from TCGA then applied CIBERSORT, a computational approach that infers immune cell content from bulk RNA-seq data. For this analysis, we assessed 6 immunogenic solid tumor types: skin cutaneous melanoma (SKCM), kidney renal clear cell carcinoma (KIRC), cervical kidney renal papillary cell carcinoma (KIRP), squamous cell carcinoma of the lung (LUSC), lung adenocarcinoma (LUAD), and bladder carcinoma (BLCA). When running CIBERSORT, we used the LM22 signature matrix designed by Newman et al[30] to estimate the relative fraction of 22 immune cell types within input mixture samples, which include an estimation of resting and activated NK cell proportions (Fig. 3D). Correlation analysis between CTLA-4 expression and CIBERSORT cell type estimation revealed that the direction of correlation in NK cells was dependent upon the activation state (Fig. 3E, Supplemental table 3). Across several tumor types, the proportion of activated NK cells was positively correlated with CTLA-4 expression while the proportion of resting NK cells was negatively correlated. CTLA-4 expression was negatively correlated with estimated proportions of resting NK cells in SKCM (p < 1 x 10^−4^), BLCA (p < 1 x 10^−3^), LUSC (p < 1 x 10^−2^), KIRP (p < 1 x 10^−2^), and KIRC (p < 1 x 10^−9^). On the other hand, estimated proportions of activated NK cells were positively correlated with CTLA-4 expression in SKCM (p < 1 x 10^−6^), BLCA (p < 1 x 10^−2^), LUSC (p < 0.05), KIRP (p < 0.05), and KIRC (p < 1 x 10^−2^). As expected, CTLA-4 expression was also positively correlated with the estimated proportions of regulatory T cells (Tregs) in each tumor type (Supplemental Table 3).

### Preclinical NK cell activation signature is associated with ipilimumab response in metastatic melanoma

While informative, bulk RNA-seq cannot resolve cell type-specific changes in gene expression. Therefore, to further investigate the relevance of the NK cell activation signature (pattern 7) to immunotherapy responses, we used our transfer learning method (projectR), to project two independent scRNA-seq datasets of ICI-treated metastatic melanoma patients[5,6] onto the 21 mouse patterns identified by CoGAPS. First, we analyzed a scRNA-seq dataset of ∼16,000 immune cells isolated from melanoma metastases. Patients in this study were treated with anti-PD-1, anti-CTLA-4, or combination anti-PD-1 and anti-CTLA-4 antibodies, and the biopsies were taken either prior to or during treatment[5]. Using the projected weights of each signature and treatment outcomes, we evaluated the association of each pattern with therapeutic response in humans. In pre-treatment biopsies, the NK cell activation signature was significantly higher in anti-CTLA-4 responsive tumors compared to non-responsive tumors (p < 1 x 10^−15^, Supplemental Fig. 3A). This is consistent with our initial finding that NK cell activation was enriched in mouse tumors treated with anti-CTLA-4.

To further examine this relationship, we tested for enrichment of the NK cell activation signature specifically in the NK cells in this dataset. While NK cells were not annotated in the study that produced this data[5], we observed that cells expressing key NK marker genes were intermixed with T cells in the lymphocyte cluster (Supplemental Fig. 3B). This is consistent with previous scRNA-seq studies that have identified subpopulations of T cells that express transcripts linked to the cytotoxic function of NK cells, such as NKT cells[31,32]. Thus, to eliminate T and NKT cells from the analysis, we performed a gene expression gating strategy that required the expression of several transcripts related to NK cell function (*NCR1*, *NKG7*, and *FCGR3A*) and a lack of the T cell transcripts (*CD4*, *CD3D*, and *CD3G)*. Gating for NK cells confirmed that the NK cell activation signature was enriched in intratumoral NK cells isolated from anti-CTLA-4 responsive tumors (Fig. 4A, p < 1 x 10^−8^). Because cells were obtained from tumor biopsies prior to the administration of anti-CTLA-4 treatment, this finding suggests that cytotoxic NK cell infiltration could be predictive of anti-CTLA-4 response. In patients treated with anti-PD-1, there was no significant difference in the NK cell activation signature between responders and non-responders regardless of whether biopsies were taken before (Fig. 4A, p > 0.05) or during (Fig. 4B, p > 0.05) treatment. In contrast, the NK cell activation signature was significantly enriched in tumors responsive to combination anti-CTLA-4 and anti-PD-1 taken before (Fig. 4A, p < 0.05) and during (Fig. 4B, p < 0.01) treatment. Using receiver operating characteristic curve (ROC) analysis, we found that the NK cell activation signature had a moderate ability to classify anti-CTLA-4 response (Fig. 4C, AUC = 0.748), suggesting that the NK activation signature has the potential utility to predict responsiveness to anti-CTLA-4 from pre-treatment tumor biopsies. These findings indicate that the presence of active NK cells within tumors is important to the clinical usage and success of anti-CTLA-4 therapies.

**Figure 4.**
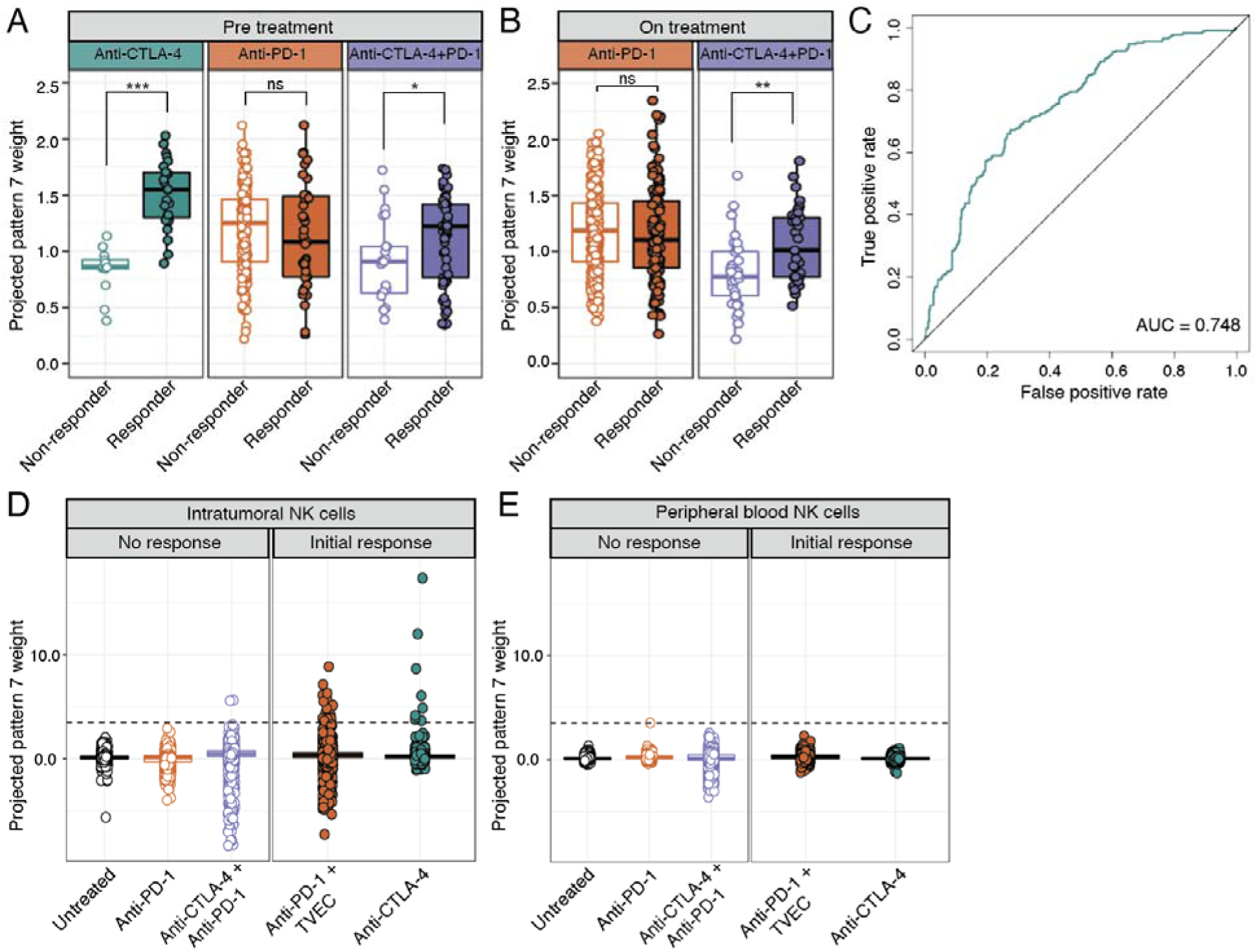
ProjectR recovers conserved immunotherapy response in intratumoral NK cells from independent human melanoma scRNA-seq datasets. **A.** Box plot of projected pattern 7 weights across intratumoral NK cells from metastatic melanoma patients prior to ICI treatment [5]. Cells are colored by therapy and separated by patient response. Increased pattern 7 is significantly associated with NK cells from patients responsive to anti-CTLA-4 or combined anti-CTLA-4 and anti-PD-1. Significant differences in mean pattern 7 weight between treatment groups are indicated by asterisks where p-values < 0.05 = *, < 0.01 = **, and p-values < 0.001 = ***. **B.** Box plot of projected pattern 7 weights across intratumoral NK cells from metastatic melanoma patients after treatment with ICI. Cells are colored by therapy and separated by patient response. Increased pattern 7 is associated with NK cells from patients responsive to combination anti-CTLA-4 + anti-PD-1. Significant differences in mean pattern 7 weight between treatment groups are indicated by asterisks where p-values < 0.05 = *, < 0.01 = **, and p-values < 0.001 = ***. **C.** ROC curve for the performance of pattern 7 weights in predicting response to anti-CTLA-4 prior to the administration of treatment. **D.** Box plot of projected pattern 7 weights across flow-sorted intratumoral NK cells from metastatic melanoma tumors that were unresponsive ICI (intrinsic resistance) or developed acquired resistance after a period of initial response[6]. The dashed line indicates the average maximum value for pattern 7 across treatment groups. NK cells with elevated pattern 7 weights are seen in patients that had an initial response to ICI, with the highest observed weights from a patient that responded to anti-CTLA-4. **E.** Box plot of projected pattern 7 weights across NK cells isolated from peripheral blood of metastatic melanoma patients that had no response to ICI (intrinsic resistance) or developed acquired resistance after a period of initial response. The dashed line indicates the average maximum value for pattern 7 from intratumoral NK cells across treatment groups. Elevated pattern 7 weights are not detected in circulating NK cells, regardless of response.

Although ICI therapy can lead to durable responses in patients with metastatic melanoma, intrinsic and acquired resistance remain major causes of mortality[33]. To examine the relationship between NK cell activation and mechanisms of therapeutic resistance, we next projected the transcriptional patterns into a dataset of NK cells isolated from melanoma metastases and matched blood samples of patients that had progressed after immunotherapy[6]. This dataset included two patients that had an initial response to ICI (acquired resistance), two patients that failed to respond to ICI (intrinsic resistance), and one patient that was not given ICI (untreated). We found high levels of the NK cell activation signature in a subset of intratumoral NK cells from the two patients who had an initial response to ICI (Fig. 4D). Consistent with our results which indicate that the NK cell activation signature is enriched in anti-CTLA-4 responsive tumors, the highest levels of the NK cell activation signature were found in NK cells from the patient responsive to anti-CTLA-4 (ipilimumab). Elevated NK cell activation signature was also found in the patient responsive to combination treatment with anti-PD-1 and oncolytic virus (pembrolizumab + TVEC). Notably, these observations were specific to intratumoral NK cells, as the NK cell activation signature was detected only at very low levels in NK cells isolated from matched peripheral blood samples (Fig. 4E). This result indicates that anti-CTLA-4 treatment leads to NK cell activation specifically within the tumor microenvironment in humans, which is consistent with observations in mice[23].

### Human NK cells express CTLA-4, which is bound by ipilimumab

CTLA-4 is a major regulator of T cells and there is growing evidence suggesting that CTLA-4 regulates other human immune cell types, including B cells[34,35], monocytes[36], and dendritic cells[37]. The role of CTLA-4 in NK cells, however, remains controversial, and the majority of the literature suggests human NK cells do not express CTLA-4[23,38–40]. However, the robust activation of intratumoral NK cells in response to anti-CTLA-4 treatment suggests that CTLA-4 may function as an NK cell immune checkpoint—similar to its role in T cells. To investigate this possibility, we first assessed the expression of *CTLA-4* transcripts in NK cells from scRNA-seq data. Indeed, some intratumoral NK cells in mice and humans express *CTLA-4* (Fig. 5A). Importantly, if the expression of *CTLA-4* is low to moderate in these NK cells, the transcripts could suffer from poor capture efficiency during scRNA-seq[41]. These technical limitations could result in the observed detection in only a handful of NK cells and require the use of *in vitro* techniques to confirm. To further investigate and validate the NK activation transcriptional signature we observed computationally, we turned to molecular biology

**Figure 5.**
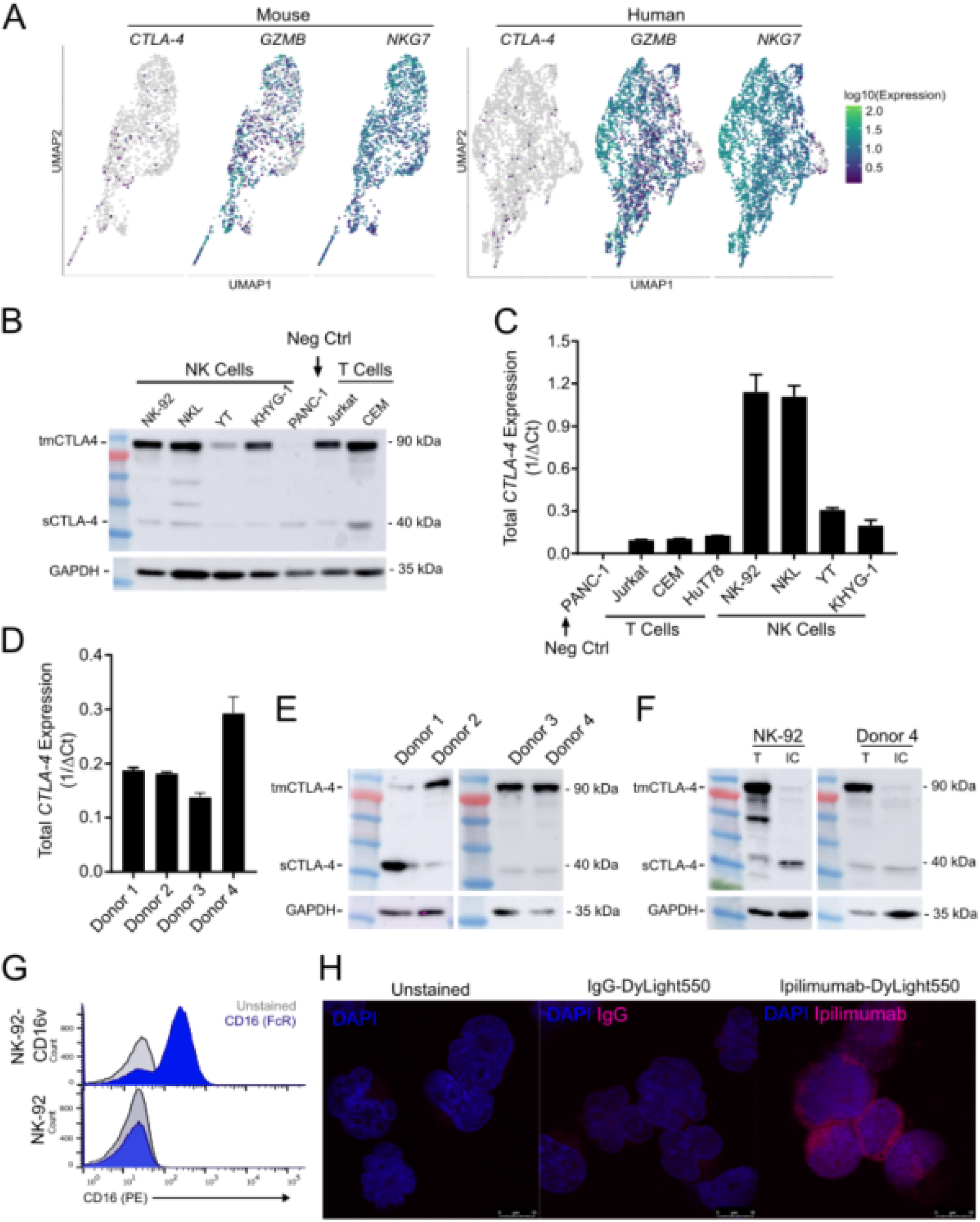
CTLA-4 is expressed by both human NK cell lines and healthy human donor-derived NK cells. **A.** UMAP dimension reduction with cells colored by single-cell gene expression for *CTLA-4* and representative immune activation genes in mouse (left) and human (right) intratumoral NK cells. The pattern of *CTLA-4* expression is consistent with the reduced ability of scRNA-seq to capture low to moderately expressed genes. **B.** Western blot demonstrating CTLA-4 expression in human NK cell lines. **C.** Quantitative real-time PCR (qRT-PCR) analysis of total *CTLA-4* expression (both isoforms) in a CTLA-4 null line (PANC-1), T cell lines (Jurkat, CEM, HuT78), NK cell lines. **D.** qrt-PCR demonstrating *CTLA-4* expression in CD56+ selected *ex vivo* unstimulated NK cells derived from healthy human donors. **E.** Western blot of CTLA-4 expression in CD56+ selected *ex vivo* unstimulated NK cells derived from healthy human donors. **F.** Western blot of total protein (T) and intracellular (IC) protein isolated from human NK cell line NK-92 and unstimulated primary human NK cells using cell surface protein biotinylation for exclusion of surface proteins demonstrating surface expression of CTLA-4 dimers and intracellular expression of CTLA-4 monomers. **G.** Flow cytometry demonstrating NK-92 does not express antibody receptor CD16. Positive control was the NK-92 line that had been transfected with a CD16 expressing plasmid, NK-92-CD16v. **H.** Immunofluorescent images of NK-92 cells stained with Dylight550-labelled ipilimumab demonstrating that ipilimumab binds to NK cell surface. Blue staining indicates DAPI. Shown are representative images of a single field of view taken via confocal microscopy (magnification, 63X, zoom, 3X).

In T cells, CTLA-4 competes with co-stimulatory receptor CD28 for B7 ligands. When CTLA-4 outcompetes CD28 for B7 binding, it prevents CD28 co-stimulatory signaling and instead provides inhibitory signaling. Anti-CTLA-4 treatment results in T cell activation by inhibiting the inhibitor; by blocking CTLA-4-B7 interactions and promoting CD28-B7 interactions. To determine if CTLA-4 could be functioning similarly in NK cells, we tested NK cells for CD28 and CD28H expression. Consistent with previous reports, we found that some NK cell lines and donor NK cells expressed CD28 and CD28H[42] by flow cytometry and qRT-PCR (Supp. Fig. 5). Thus, human NK cells appear to express both CTLA-4 and CD28, supporting a parallel role for these receptors in T cells and NK cells.

### Ipilimumab binds to CTLA-4 expressed on the NK cell surface independent of CD16

We next wanted to determine if the anti-CTLA-4 antibody, ipilimumab, was capable of binding to CTLA-4 expressed on the NK cell surface. To do so, we fluorescently labelled anti-CTLA-4 (Ipilimumab) to probe for ipilimumab binding to the NK cell surface by immunofluorescence microscopy. One potential complication is nonspecific binding of ipilimumab to NK cells. Human NK cells express antibody receptors (e.g., Fc receptor CD16) which can bind to the constant region of an antibody regardless of the antibody’s specificity. [43]. To exclude the possibility of nonspecific ipilimumab-NK cell interactions, we used the human NK cell line NK-92, which lacks generic antibody receptors (i.e., CD16) (Fig. 5G). Immunofluorescence imaging demonstrated that fluorescently labeled anti-CTLA-4, but not the IgG control, was capable of binding to NK-92 through recognition of CTLA-4 on the cell surface (Fig. 5H). The specificity of the stain was confirmed using the CTLA-4 null line PANC-1 (Supplemental Fig. 4E. We saw abundant surface expression of CTLA-4 by immunofluorescence confirming the results shown in Figure 5F. To the best of our knowledge, this is the first demonstration that anti-CTLA-4 (ipilimumab) can directly interact with human NK cells via a CD16-independent mechanism.

### NK cell activation regulates CTLA-4 expression

CTLA-4 expression is modulated in response to T cell activation via CD28 and T cell receptor signaling[44]. To investigate if *in vitro* NK cell activation would similarly modify CTLA-4 expression in NK cells, we exposed NK cells to a variety of cytokines (IL-2, IL-12, IL-15, L-18) that activate NK cells and alter NK cell expression of other immune checkpoints (i.e. PD-1)[45][46] (Fig. 6A). Human NK cells, with the exception of NK cell line NK-92, had a drastic reduction in CTLA-4 after 24-hour exposure to IL-2. IL-15 also caused a reduction in CTLA-4 expression in all NK cells tested except NKL. Alternatively, IL-12 and IL-18 increased CTLA-4 expression in a subset of NK cell lines, including primary donor NK cells.

**Figure 6.**
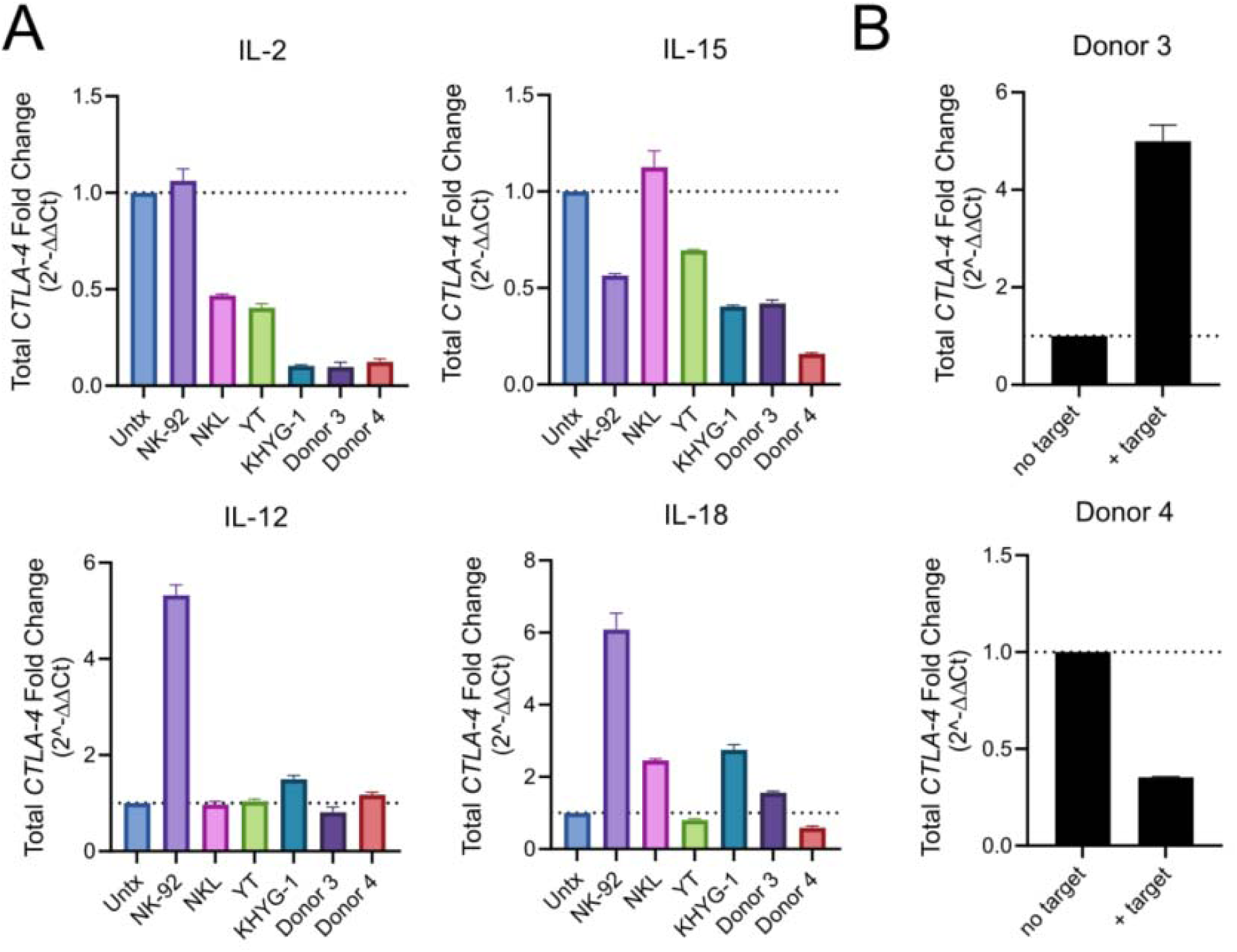
NK cell activation regulates CTLA-4 expression. **A.** Effect of 24 hour stimulation with IL-2, IL-12, IL-15 and IL-18 on NK cell CTLA-4 expression as determined by qRT-PCR. **B.** Effect of target cell exposure (K562-4-1BB-mbIL-21) on NK cell CTLA-4 expression as determined by qRT-PCR.

Target cell recognition is another means to activate NK cells. Since cytokine-activated and target-cell activated NK cells have distinct transcriptional phenotypes[47] we also investigated target cell-mediated NK cell activation on NK cell CTLA-4 expression by exposing NK cells to engineered target cells (K562-4-1BB-mbIL-21 cells) (Fig. 6B). Although we saw divergent responses in the primary NK cells from two donors, target cell exposure clearly modulated CTLA-4 expression. These data demonstrate that although responses are variable, human NK cell activation, via cytokine and target-cell stimulation, alters NK cell expression of CTLA-4. Combined with the observation that anti-CTLA-4 antibodies bind human NK cells, these results suggest CTLA-4 may be an NK cell checkpoint and drive the computationally identified signature of NK cell activation in anti-CTLA-4 responsive tumors. Taken together, these results confirm the utility of CoGAPS and projectR to identify conserved biological processes between preclinical models and human patients that contribute to clinical outcomes.

## 3. Discussion

Using a combination of state-of-the-art computational algorithms and molecular biology approaches we demonstrate that transfer learning can be used to elucidate complex tumor immune responses to immunotherapy that are conserved between species. Specifically, in this study we demonstrate that the Bayesian non-negative matrix factorization approach CoGAPS associates intratumoral NK cell activation and anti-CTLA-4 response. These findings extend work done by Gubin et al.[4], which used scRNA-seq and CyTOF to profile changes in immune cells from mouse sarcomas following immunotherapy treatment. Using standard scRNA-seq analysis methods, Gubin et. al. did not detect NK cell activation from the scRNA-seq data in anti-CTLA-4 treated tumors, however their subsequent CyTOF analysis revealed prominent upregulation of NK cell granzyme expression specific to anti-CTLA-4 treatment[4]. To bridge these datasets, we used our transfer learning method, projectR, to demonstrate that the scRNA-seq signature of NK cell activation in response to anti-CTLA-4 therapy was preserved at the protein level in the paired CyTOF data (Supplemental Figure 1B). Using several additional clinical datasets, we determine a robust association with this signature and anti-CTLA-4 activity in human tumors.

The likely source of difference in the features identified in our study and Gubin et al.[4] is the analysis method for cell state identification. Notably, Gubin et al.[4] employed clustering for their analysis, which is a central step in standard scRNA-seq pipelines to group discrete cell populations that share similar transcriptional profiles. In addition to cell type specific activities, transcriptional profiles simultaneously include cellular processes such as activation, exhaustion, and cell signaling, which are not necessarily captured by clustering approaches. These processes are particularly important in the context of immune cells within the tumor microenvironment, where cells may undergo stimulation or dysregulation. In contrast to clustering, matrix factorization methods can distinguish the molecular dynamics present in a dataset in an unsupervised manner. Our matrix factorization method, CoGAPS, was able to identify NK cell activation in response to treatment directly—without the need for clustering, differential expression analyses, or additional technologies— highlighting the advantage of our approach compared to standard analysis methods. Therefore, CoGAPS is able to identify immunotherapy induced cellular changes that may be missed by alternative methods.

Cross-species analysis is complicated by biological and technical factors, including batch effects due to experimental platforms and species-specific differences. Attempts to integrate single-cell data from mice and humans often rely on batch correction methods, which adjust the expression levels of genes within cells from each species to resemble each other[48,49]. In contrast, transfer learning takes low-dimensional gene expression signatures learned from latent space techniques (e.g, CoGAPS) on one dataset and maps them to another—without the need for batch correction. Previous transfer learning studies have demonstrated the ability to transfer immune cell type labels between datasets[50] and in cross-species analysis of developmental processes[7,9]. Here, we sought to further test the ability of transfer learning to elucidate complex tumor immune responses to immunotherapy that are preserved between preclinical models and human tumors. Thus, after identifying NK cell activation in mouse tumors treated with anti-CTLA-4, we used projectR to probe human datasets for an association with clinical outcomes. Despite known differences between mice and human NK cell surface receptors[51], our approach was able to analyze homologous genes and confirmed that the NK cell activation signature we observed in anti-CTLA-4 treated mice was conserved between species and cancer types. In addition to being a conserved response to anti-CTLA-4, we found that elevated levels of the NK cell activation signature was associated with better clinical response to anti-CTLA-4 treatment. This supports the continued observation that key biological roles of NK cells are shared between species[51]. Computationally, this supports that NK cell state transitions learned with CoGAPS are also preserved across species and between datasets from across technical batches. While we focus this study on CoGAPS analysis, we note that the transfer learning approach can be combined with other latent space techniques to identify gene expression signatures from single-cell data (eg. PCA, clustering, and other forms of linear matrix factorization) to identify the preservation of additional features learned from alternative approaches[7,24]. Future extensions to this approach are needed to enable transfer learning from emerging non-linear methods for inference of more complex cell state transitions and gene regulatory networks.

In this study, we have concentrated our cross-species transfer learning analysis analysis and experimental validation on the NK cell activation signature identified by CoGAPS (pattern 7). We chose pattern 7 as an interesting case of computationally identified biology for several reasons: (1) pattern 7 was the most clearly associated with a specific cell type and treatment; (2) increased expression of NK cell activation markers had been noted in anti-CTLA-4 treated mice from the original CyTOF analysis[4]; (3) there is growing evidence that CTLA-4 is expressed by non-T cell human immune cell types[34–37]; and (4) recent work found that human NK cells express PD-1 and are modulated by anti-PD-1 therapy[52,53]. Therefore, we hypothesized that CTLA-4 was similarly expressed by human NK cells and activated by anti-CTLA-4 antibodies. In agreement with our findings, several reports highlight an interesting relationship between NK cells and anti-CTLA-4 response in humans. In melanoma patients treated with anti-CTLA-4, a higher percentage of circulating mature NK cells is correlated with improved overall survival, and NK cells isolated from responsive patients have increased cytolytic activity compared to NK cells isolated from non-responders[54]. In B16 melanoma models, NK cells and CD8+ T cells synergistically clear tumors in response to anti-CTLA-4 and IL-2 treatment[55]. Furthermore, anti-CTLA-4 has been shown to increase transcriptional markers of NK cell cytotoxic activity in CT26 colon carcinoma tumors[23]. While future mechanistic studies are needed to fully elucidate the specific function(s) of CTLA-4 in NK cell biology, these findings support the computational approaches employed in this study.

We leverage our computational findings to guide molecular experiments, through which we provide a rationale for NK cell activation in response to anti-CTLA-4 by demonstrating that NK cells constitutively express CTLA-4 on their cell surfaces and bind anti-CTLA-4 antibodies. A number of immune checkpoints are expressed by both T cells and NK cells. For example, recent studies have found that NK cells within several human and mouse tumor types express PD-1, and that ligands for these checkpoint receptors negatively regulate NK cell activity[53,56]. Consistent with this, blocking PD-1 receptors with anti-PD-1 therapy enhances NK cell-mediated anti-tumor responses[22], and NK cell infiltration correlates with clinical responsiveness to anti-PD-1 therapy[57]. Despite growing evidence for the role of checkpoint receptors in NK mediated anti-tumor responses, the expression of CTLA-4 by NK cells has been disputed in the literature. While mouse NK cells inducibly express CTLA-4 in response to IL-2[45], a recent study was unable to detect CTLA-4 on the surface of intratumoral murine NK cells[23]. An earlier study in humans also reported that NK cells from healthy donors do not express CTLA-4 [38]. Contrary to these earlier reports, our results demonstrate CTLA-4 is constitutively expressed by circulating healthy human donor NK cells and human NK cell lines. One possible explanation for why previous studies have failed to identify the expression of CTLA-4 by human NK cells is the reliance on flow cytometry in these studies. Flow cytometry can be limited by challenges related to the generation of antibodies and further complicated by the rapid surface expression dynamics of CTLA-4[58]. In support of this explanation, we too fail to detect intracellular or surface CTLA-4 expression when using flow cytometry (Supplemental Fig 4A and B)), even though we are able to unequivocally demonstrate CTLA-4 expression at the RNA and protein level by qRT-PCR and western blot in *ex vivo* unstimulated healthy donor NK cells (Fig. 5B-E), as well as surface expression using immunofluorescence and biotinylation (Fig. 5G). Consistent with previous studies[59,60], we show that human NK cells express CD28 and CD28H (Supplemental Fig. 5), a co-stimulatory receptor that competes with CTLA-4 for the binding of B7 ligands. The expression of B7 on tumor cells also enhances NK recognition and lysis of tumors through CD28-B7 interactions[59–65]. In addition, we show that CTLA-4 expression by human NK cells cultured *in vitro* is modulated in response to NK cell activation (Figure 6). These findings suggest that CTLA-4 may have similar functions in NK cells and effector T cells[44].

In addition to informing molecular experiments, transfer learning to human cohorts with clinical outcomes can facilitate translational research in developing mechanistic biomarkers from preclinical models. In this study, our transfer learning analysis demonstrates that the NK cell activation signature we learned in the preclinical scRNA-seq data is conserved in anti-CTLA-4 responsive human tumors prior to anti-CTLA-4 treatment. Moreover, the amount of this NK cell activation pattern prior to treatment correlates with the clinical response to anti-CTLA-4 in metastatic melanoma. This indicates that NK cells must already be activated within tumors in order to have improved tumor clearance by the addition of anti-CTLA-4. Future transfer learning analyses on large cohort studies of anti-CTLA-4 treated tumors with genomics data could further delineate its role as a potential predictive biomarker. However, these datasets are currently lacking in the literature, which limits our ability for such computationally-driven biomarker analysis. Still, our present study found that the NK cell activation signature was observed in tumors responsive to anti-CTLA-4 alone or in combination with anti-PD-1, suggesting that this signature is a specific response to therapies that include anti-CTLA-4. *In vivo*, it is possible that the on-target interaction of anti-CTLA-4 antibodies and CTLA-4, as well as Fc receptor engagement on NK cells contribute to anti-tumor activity.

In the context of therapeutic resistance, we detected NK cells with high expression of the NK cell activation signature in patients that developed acquired, but not primary, resistance to immunotherapy. This indicates that intratumoral NK cell activation is able to identify patients that have an initial response to therapy. Consistent with a relationship between NK cells and anti-CTLA-4 response, we observed the highest levels of NK cell activation in intratumoral NK cells isolated from a patient that had an initial response to anti-CTLA-4. This enrichment was absent in patients that were unresponsive to anti-PD-1, either alone or in combination with anti-CTLA-4. Surprisingly, the NK cell activation signature was also elevated in a patient that initially responded to combination anti-PD-1 and oncolytic virus. This could be due to the fact that infection of tumors with oncolytic viruses can activate NK cells and stimulate NK-mediated anti-tumor immunity[66]. Furthermore, since this observation was specific to intratumoral NK cells and not circulating NK cells, approaches to transcriptionally profile patients using peripheral blood may be limited in identifying signatures related to clinical outcomes. It will be important for future studies to determine the role of NK cells in anti-CTLA-4 response and resistance.

As scRNA-seq atlases become increasingly prevalent in cancer research, computational tools to generalize findings across species are necessary. This work describes a useful computational approach for studying cancer immunotherapy that is able to identify cellular responses preserved across different data modalities, species, and patients. This provides a powerful method for extrapolating relevant information while avoiding the unique biases of individual technologies (i.e., dropout in scRNA-seq, biased selection of genes in CyTOF, or aggregate transcriptional profiles in bulk RNA-seq). In addition, it allows the comparison of different treatment conditions, disease states, and tumor types. Therefore, we provide a framework for cross-species data analysis, with the feasibility to integrate preclinical and clinical genomics datasets. Following the integration of larger clinical cohort single-cell studies, we anticipate that these methods will aid in the prediction of patient prognosis and therapeutic response. Due to the flexibility of our approach, these algorithms can be used to study the treatment of disease in a variety of contexts. The ability to rapidly identify conserved responses to therapy between mice and humans will help bridge basic science and clinical research and advance our ability to understand and treat disease.

## 4. Methods

### Data collection

In this study, we used three public scRNA-seq datasets generated by different groups using droplet-based profiling technologies. Read counts for each scRNA-seq dataset were obtained from NCBI’s Gene Expression Omnibus.

For CoGAPS analysis on preclinical immunotherapy samples, we used a scRNA-seq dataset containing ∼15,000 flow-sorted CD45+ intratumoral cells from mouse sarcomas that were collected during treatment with either control monoclonal antibody, anti-CTLA-4, anti-PD-1, or combination anti-CTLA-4 and anti-PD-1[4]. This data was acquired with the 10x Genomics Chromium platform, using v1 chemistry. The accession number for this dataset is GSE119352. The scRNA-seq data was complemented by paired mass cytometry data stored in the FLOW Repository under FR-FCM-ZYPM. Data of 5 replicates per treatment were processed using the R package cytofkit version 0.99.0 and used for transfer learning analysis.

For transfer learning to human samples, we used two scRNA-seq datasets of intratumoral immune cells from metastatic melanoma patients. To first test the relationship between our preclinical CoGAPS patterns and clinical outcome, we used a scRNA-seq dataset containing ∼16,000 flow-sorted CD45+ intratumoral cells obtained from 48 human melanoma tumor biopsies from 32 patients at baseline or after treatment with either anti-CTLA-4, anti-PD-1, or combination anti-CTLA-4 and anti-PD-1[5]. This data was acquired with Smart-seq2. The accession number for this dataset is GSE120575.

Next, to confirm the observed relationship between our preclinical NK activation signature and response to anti-CTLA-4, we used a scRNA-seq dataset containing ∼40,000 flow-sorted NK cells from matched blood and tumor samples obtained from 5 patients with melanoma metastases[6]. Two patients had an initial response to treatment with anti-CTLA-4 or anti-PD-1 with oncolytic virus. Two patients failed to respond to combination anti-CTLA-4 and anti-PD-1 or anti-PD-1. One patient was not treated with immunotherapy. This data was acquired with the 10x Genomics Chromium platform, using v2 chemistry. The accession number for this dataset is GSE139249.

In addition, bulk RNA-seq was downloaded from The Cancer Genome Atlas[25]. In this case, level 3 RSEM normalized across 33 tumor types were accessed from the Broad Institute TCGA GDAC Firehose (http://gdac.broadinstitute.org/runs/stddata 2016_01_28/data/) and log2-transformed. CIBERSORT scores for this data were obtained from Thorsson et al.[67].

These datasets were used for pattern discovery and transfer learning as described below.

### Dimensionality reduction and cell type identification

Cell type inference analyses were performed for the Gubin et al. dataset with the standard Monocle3 workflow using package version 0.2.0. Dimensionality reduction and visualization for scRNA-seq data were performed using Uniform Manifold Approximation and Projection (UMAP)[68]. Briefly, the first 15 principal components were used as input into the reduce_dimension function. Canonical cell type marker genes as described in Gubin et al. were used to annotate cells[4].

### Mouse pattern discovery and gene set analysis using CoGAPS

CoGAPS analysis was performed using the R/Bioconductor package CoGAPS version 3.5.8 to analyze the mouse sarcoma dataset from Gubin et al.[4]. Genes with a standard deviation of zero were removed prior to analysis. The log2 transformed count matrix of remaining genes across all samples was used as input to the CoGAPS function. Default parameters were used, except nIterations = 50,000, sparseOptimization = True, nSets = 12. The input parameters for nPatterns was determined empirically, by testing over a range of dimensions. When the nPatterns input was set to 3 we obtained results that identified immune cell lineage. We reasoned that additional patterns could further identify biological processes in the data related to treatment. We initially tested 50 patterns, however, many of the patterns highlighted few cells, indicating an over-dimensionalization of the data. We obtained stable results when nPatterns was set to 25, with the final CoGAPS dataset stabilized at 21 patterns. Genes highly associated with each pattern were identified by calculating the PatternMarker statistic[18]. The CalcCoGAPSStat function was used to identify pathways significantly enriched in each pattern for the MSigDB hallmark gene sets[15] and PanCancer Immune Profiling panel from NanoString Technologies.

### Pseudotime analysis

To perform pseudotemporal ordering, the dataset was subset to relevant cell types and treatments based on the desired analysis. Due to the association between pattern 7 and activation state markers, we chose the most active terminus of the trajectory as the end state. Thus, the root node of the trajectory was assigned by identifying the region in the UMAP dimensional reduction with low CoGAPS pattern 7 weights. Pseudotime values were assigned to cells using the order_cells function from Monocle3 version 0.2.0. Genes with significant expression changes as a function of pseudotime were identified using the graph_test function, using a multiple-testing corrected q-value cutoff of 0.01.

### Linear modeling

TCGA expression and metadata were aggregated using the R/Bioconductor package TCGAbiolinks version 2.14.1[69], and was used as input for transfer learning as described below. Samples were restricted to those that were labeled as “Primary solid tumor” (n=9113), “Recurrent solid tumor” (n=46), and “Metastatic” (n=394) in the “definition” column of the TCGA metadata, which resulted in 9,553 total samples. Measures of overall survival and age at diagnosis for TCGA samples were taken from those aggregated by Liu et al.[70]. After scaling and centering the data, linear models were run according to the following equation:

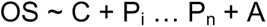

Where OS equals overall survival, C represents cancer type as a categorical variable, P_i…P_n represent each transcriptional signature, or pattern, as separate continuous covariates, and A equals age at diagnosis. Linear models fit per cancer type were run on samples belonging to each respective cancer, and did not include the cancer type covariate C from the equation above. Models looking at the relationship between age and patterns replaced OS in the equation above with A.

### Survival analysis

Kaplan Meyer plots were generated in R using the survfit function from the survival package version 3.1-12, and the ggsurvplot function from the survminer package version 0.4.6. Samples were split into those in the top 5% of pattern 7 scores, and those in the bottom 95% (i.e. all other samples).

### Correlation analysis

To compare the expression of CTLA-4 and CIBERSORT scores for various immune cell types across immunogenic solid tumors from TCGA, we calculated the Spearman correlation coefficients using the cor.test function in R.

### Transfer learning

To examine whether the mouse patterns corresponded to similar immunotherapy responses in human data, we used The R/Bioconductor package projectR[24] version 1.0.0 to project the expression matrix from several datasets into the CoGAPS pattern amplitude matrix[7]. The CoGAPS result object and the expression matrix from a human dataset is used as input to the projectR function. This algorithm returns a new pattern matrix, which estimates the role of each pattern in each cell of the human dataset. This comparison of pattern across species usage enabled us to determine how each pattern defines features present in the human dataset (i.e. cell types and immune cell activation). Homologous genes present in the mouse and human data were retained for projection. Genes without homologs in the human data were removed.

### Pattern performance of predicting anti-CTLA-4 response

The projected pattern weight is a continuous range of values, instead of a binary outcome. Using the individual projected pattern weight for each cell and a binary response outcome to anti-CTLA-4, we performed ROC curve analysis using the ROCR package, version 1.0-7 to determine the true-positive rates versus false-positive rates of pattern 7 weights to classify response. The area under the ROC curve was used as the quality metric to determine the prediction performance.

### Cell lines and materials

All human NK cell lines (NK-92, NK-92-CD16v, NKL, YT and KHYG-1) were kindly provided by Dr. Kerry S. Campbell (Fox Chase Cancer Center, Philadelphia, PA). The NK-92-CD16v expressed GFP due to transduction with pBMN-IRES-EGFP containing the Fc RIIIA construct. All NK cell lines were cultured as previously described[71]. Fresh healthy donor NK cells were purchased from AllCells (PB012-P). These NK cells were positively selected from donor peripheral blood using CD56 positivity. Donor NK cell purity was 98-99%. Donor 3 and Donor 4 were expanded using engineered antigen presenting cells (K562-4-1BB-mbIL-21) according to the protocol[72]. CTLA-4 overexpressing Jurkat cell line was generated using lentiviral transduction purchased from G&P BIosciences (Product ID: LYV-CTLA4, SKU#: LTV0710) which contained full length human CTLA-4 gene subcloned into lentiviral expression vector pLTC with an upstream CMV promoter with puromycin selection marker. Jurkat cells were transduced using millipore sigma’s spinoculation protocol. In brief, lentiviral particle solution was added to 2 X 10^6^ Jurkat cells at a final multiplicity of infection of 1, 5 and 10. Cells were centrifuged at 800 xg for 30 minutes at 32°C then resuspended in complete growth medium for 3 days. After three days, cells were resuspended in complete medium containing 5 ug/mL puromycin overnight for selection. Selection was performed twice.

### qRT-PCR

RNA was isolated using the PureLink RNA Mini Kit (Ambion). The RNA concentration was measured using NanoDrop 8000 (Thermo Fisher Scientific). cDNA was generated from 20-100 ng of RNA using the GoTaq 2-step RT-qPCR System (Promega). qPCR was performed with SYBR Green on a StepOnePlus real-time PCR system (Applied Biosystems). Gene expression was normalized to HPRT and analyzed using 1/DCt method with triplicates. RNA was isolated using the PureLink RNA Mini Kit (Ambion). The RNA concentration was measured using NanoDrop 8000 (Thermo Fisher Scientific). cDNA was generated from 20-100 ng of RNA using the GoTaq 2-step RT-qPCR System (Promega). qPCR was performed with SYBR Green on a StepOnePlus real-time PCR system (Applied Biosystems). Gene expression was normalized to HPRT and analyzed using 1/DCt method with triplicates.

Primers used were:

CTLA-4: (F: CATGATGGGGAATGAGTTGACC; R: TCAGTCCTTGGATAGTGAGGTTC)

CD28: (F: CTATTTCCCGGACCTTCTAAGCC; R: GCGGGGAGTCATGTTCATGTA)

CD28H: (F:CCCTGCAAGAAGCCTCAAG; R: CCTTTGTCCACTTAACACGGAG)

HPRT: (F: GATTAGCGATGATGAACCAGGTT; R: CCTCCCATCTCCTTCATGACA)

### Western Blot

Cells were lysed in boiling buffer with EDTA (Boston BioProducts) supplemented with 1X protease and 1% phosphatase inhibitor prepared following the manufacturer’s protocols (Sigma-aldrich, Cat.No. 11697498001 and P5726). Cleared lysate concentrations were obtained by a DC Protein Assay (BioRad). Lysates 30-50 ug were run on SDS-PAGE gels and transferred to nitrocellulose membranes (GE Healthcare). Western blots were conducted using anti-CTLA-4/CD152 (LS C193047, LSbio) at concentrations of 1:1000 diluted in 5% milk in PBST. Secondary antibody was anti-rabbit IgG, HRP linked (Cell Signaling) used at 1:1000. Chemiluminescent substrate (Pierce) was used for visualization.

### Flow Cytometry

All cells were aliquoted into Eppendorf tubes, spun at 5000 rpm for 1 minute at 4°C, washed twice with HBSS (Fisher Scientific Cat. No. SH3058801), and resuspended in 50 µL of FACS buffer (PBS plus 1% BSA) and blocked with µL 1 human Fc block (BD Biosciences, 564219) for 20 minutes at 4°C. Labeled antibodies were then added at the manufacturer’s recommended concentrations and incubated at 4° C for 30 minutes, with vortexing at 15 minutes. Cells were then washed with FACS buffer twice and resuspended in FACS buffer or fixative (1% PFA in PBS). Flow antibodies included anti-human CD152 (CTLA-4) (BD Bioscience 555853), CD28 (Biolegend 302907), and CD28H (R&D Systems, cat#MAB83162). The CD152 antibody has previously been shown to adequately detect CTLA-4 expression on both human T and B cells (29). Samples were run in the Georgetown Lombardi Comprehensive Cancer Center Flow Cytometry & Cell Sorting Shared Resource using BD LSRFortessa. Analyses were performed using FlowJo (v10.4.1).

### Immunofluorescence

Ipilimumab was acquired from the Medstar Georgetown University Hospital. Ipilimumab was labelled with Dylight550 fluorophore using the Dylight550 Conjugation Kit (Fast)-Lightning-Link (abcam, ab201800). In short, Ipilimumab was diluted from 5 mg/mL to 2 mg/mL using sterile PBS. Human IgG (Jackson ImmunoResearch, 009-000-003) was diluted from 11mg/mL to 2 mg/mL using sterile PBS. 1 uL of modifying reagent was added to 10 uL diluted ipilimumab and 10 uL diluted human IgG. 10 uL antibody was then added to the conjugation mix and incubated at room temperature in the dark for approximately 6 hours. 1 uL of quencher reagent was added to the labeled ipilimumab and the antibody was stored in the dark at 4°C. NK-92 and PANC-1 cells were collected and washed with cold PBS and brought to a final concentration of 1 X 10^6^ cells/mL in staining buffer (1% BSA in PBS) in 50 uL. 50 uL of labelled ipilimumab or human IgG was added to cells to yield a final concentration of 1 ug/mL antibody. Cells were incubated in the dark at 4°C for 1 hour. After incubation, cells were pelleted and washed three times with cold PBS. Cells were brought to a final concentration of 0.5 X 10^6^ cells/mL and 100 uL was immobilized on slides using cytospin (Cytospin 2, Shandon) for 5 mins at 1000 rpm. Following immobilization cells were fixed with 4% PFA for 10 minutes at room temperature then washed three times with cold PBS. Coverslips were mounted using VectraSheild mounting media with DAPI and sealed using clear nailpolish and allowed to dry overnight in the dark. Analyses were performed with the Leica SP8 AOBS laser scanning confocal microscope.

### Cell Surface Biotinylation

Cell surface biotinylation of NK92, NKL, YT and KHYG-1 cells was performed with the Pierce Cell Surface Protein Isolation kit (Thermo Scientific, cat#89881) according to the manufacturer’s protocol. In brief, 4×10^8^ cells were pelleted and washed with cold PBS then incubated with EZ-LINK Sulfo-NHS-SS-biotin for 30 min at 4°C followed by the addition of a quenching solution. Another 1X10^6^ cells were collected and saved for total cell western blotting. Cells were lysed with lysis buffer (500 μL) containing the cOmplete protease inhibitor cocktail (Roche, cat#11697498001). The biotinylated surface proteins were excluded with NeutrAvidin agarose gel (Pierce, 39001). Samples were diluted 50 ug in ultrapure water supplemented with 50 mM DTT. Lysates were subjected to Western blotting with the anti-CTLA-4 antibody described above.

### NK cell stimulation

Cell lines or expanded primary NK cells were stimulated with 100 U/mL IL-2 (NCI preclinical repository), 5 ng/mL IL-12 (R&D Systems, cat#219-IL-005), 10 ng/mL IL-15 (NCI preclinical repository), 50 ng/mL IL-18 (Invitrogen, cat#rcyec-hil18) or 500 U/mL IFNg (Sigma Aldrich, cat#I3265) for 24 hours. Cell pellets were collected and processed for rt-qPCR as described above. Cell lines or expanded primary NK cells were stimulated with 3 ug/mL CD28 activating antibody (Biolegend, cat#302933) for 24 hours.

## Data availability

| Resource | Source | Identifier |
| --- | --- | --- |
| Mouse sarcoma ICI therapy CD45+ scRNA-seq | Gubin et al. | GSE119352 |
| Mouse sarcoma ICI therapy CyTOF | Gubin et al. | FR-FCM-ZYPM |
| Treatment-naive biopsies from 33 tumor types bulk RNA-seq | TCGA |  |
| Human metastatic melanoma CD45+ cells taken pre- and post-ICI scRNA-seq | Sade-Feldman et al. | GSE120575 |
| Human metastatic melanoma with resistance post-ICI therapy flow sorted NK cells scRNA-seq | de Andrade et al. | GSE139249 |

## Code availability

All code used for the analysis is available at: https://github.com/edavis71/projectR_ICI

## Acknowledgments

This work was supported by grants from the NIH (R01CA177669, U01CA196390, and U01CA212007 to EJF) and AWS cloud credits for education. EFDM is supported through NIH F31 CA250135-01A1. EMJ acknowledges funding from the Broccoli Foundation, The Bloomberg∼Kimmel Institute for Cancer Immunotherapy, and The Skip Viragh Center for Pancreas Cancer Clinical Research and Patient Care, and The Commonwealth Foundation for Cancer Research. EMJ is also supported through NIH R01CA184926, as well as Stand Up To Cancer which is a program of the Entertainment Industry Foundation administered by the American Association for Cancer Research. EJF and EMJ are also supported through the NIH P50CA062924 and P30CA006973, the Lustgarten Foundation, the Allegheny Health Network, and the Emerson Foundation (640183). LVD is supported through P30CA006973 and R50CA243627. LMW is also supported through NIH CA50633 and CA51008. AAF is supported through NIH F30 CA239441.

We thank Dr. Kerry Campbell for providing the NK cell lines, Dr. Dean Lee for providing the NK cell expansion feeder cells, and Dr. Todd Armstrong, Dr. Kerry Campbell, Dr. Luciane Kagohara, Dr. Sandra Jablonski, Dr. Michael Atkins and Dr. Jackie Zimmerman for feedback on the manuscript.

The graphical abstract and Figure 1A were created using Biorender.com.

## Supplemental Data

**Supplemental Figure 1:**
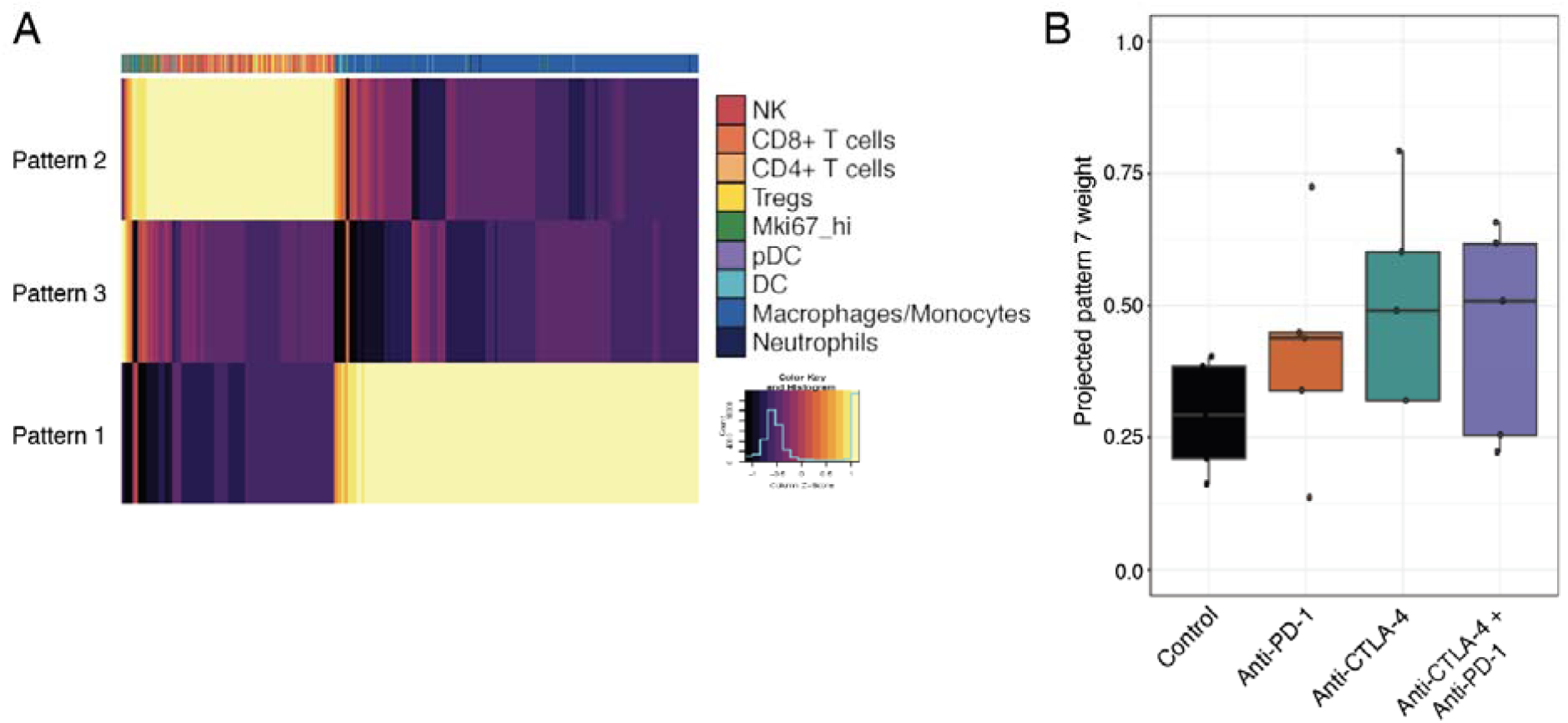
CoGAPS patterns identify immune cell lineage and transfer across data modalities. A. Heatmap of transcriptional signatures (patterns) identified with CoGAPS. When CoGAPS is performed at low dimensionality, here being 3 patterns, the identified transcriptional signatures segregate cells by immune cell lineage. Pattern 3 is relatively flat across all cells, while patterns 1 and 2 define myeloid and lymphoid lineage cells, respectively. B. Boxplot of the projected NK cell activation signature (pattern 7) weights in tumor infiltrates from mouse tumors analyzed by mass cytometry on day 11 after treatment. Each point represents a replicate sample. For each replicate, the mean protein expression of 37 genes was used as input for projectR. The NK cell activation signature is highest in lymphocyte samples treated with anti-CTLA-4, either alone or in combination with anti-PD-1.

**Supplemental Figure 2:**
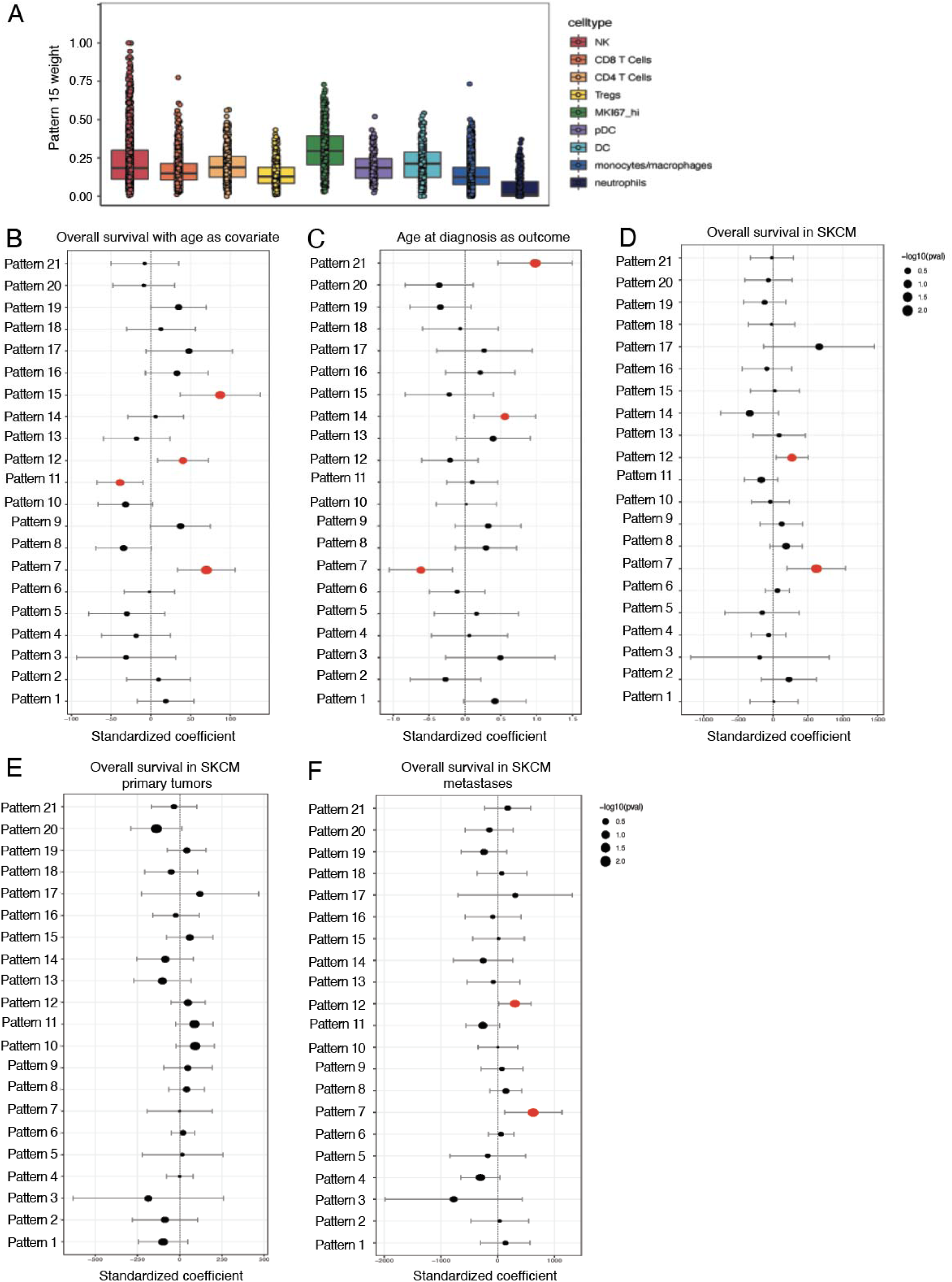
Effect sizes of pattern associations with TCGA tumor survival. A. Boxplot of pattern 15 weights across each immune cell type from mouse sarcomas. Cells with the highest pattern 15 weights are observed in NK cells and Mki67hi proliferative lymphocytes. B. The output is shown from a multiple linear regression model that predicts overall tumor survival from the CoGAPS transcriptional patterns, while also adjusting for cancer type and patient age as covariates. Standardized coefficients (i.e. data was scaled and centered) representing the strength and direction of association for each pattern are shown on the x-axis, with error bars representing coefficient +− 1.96 * standard error, and point size scaled to the coefficient’s p-value. Patterns 7 and 15 are most strongly positively associated with overall survival, with pattern 7 being most significantly positively associated (p < 2.7 x 10^−4^). C. The output is shown from a multiple linear regression model that predicts age of diagnosis from the CoGAPS transcriptional patterns, while also adjusting for cancer type as a covariate. (p < 0.017) D. The output is shown from a multiple linear regression model that predicts overall tumor survival in SKCM from the CoGAPS transcriptional patterns, while also adjusting for patient age as a covariate. Pattern 7 is the most significantly positively associated with overall survival in SKCM (p < 0.005). E. The output is shown from a multiple linear regression model that predicts overall tumor survival in SKCM primary tumors from the CoGAPS transcriptional patterns, while also adjusting for patient age as a covariate. Pattern 7 is not associated with overall survival in primary SKCM (p > 0.05). F. The output is shown from a multiple linear regression model that predicts overall tumor survival in SKCM metastases from the CoGAPS transcriptional patterns, while also adjusting for patient age as a covariate. Pattern 7 is the most significantly positively associated with overall survival in SKCM metastases (p < 0.016).

**Supplemental Figure 3:**
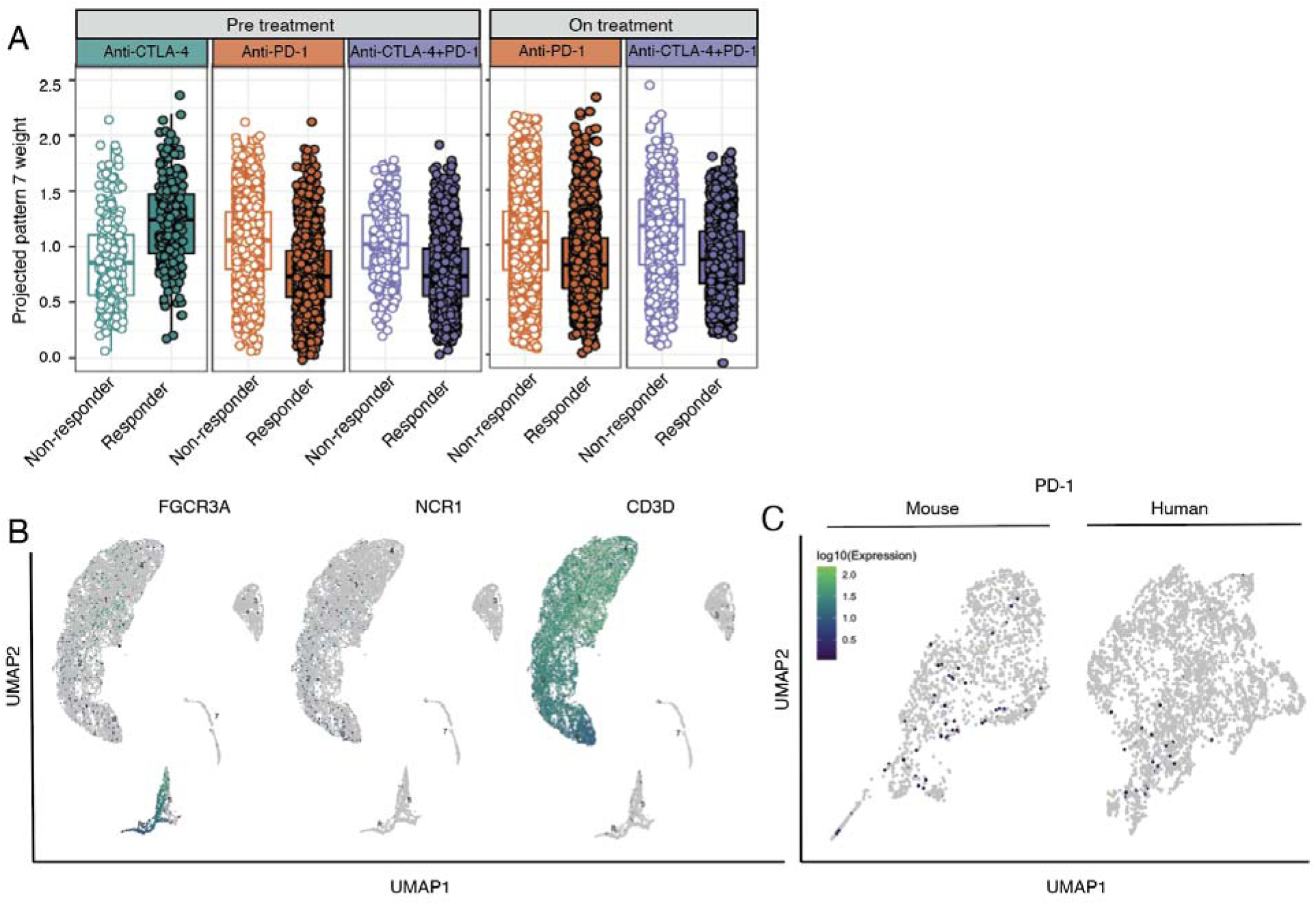
NK cell activation signature is associated with anti-CTLA-4 response. A. Box plot of projected pattern 7 weights across intratumoral immune cells from metastatic melanoma patients prior to ICI treatment [5]. Cells are colored by therapy and separated by patient response. Increased pattern 7 is associated with immune cells from patients responsive to anti-CTLA-4. B. UMAP dimension reduction with cells colored by single-cell gene expression for representative NK and T cell marker genes. C. UMAP dimension reduction with cells colored by single-cell gene expression for PD-1 in mouse (left) and human (right) intratumoral NK cells. Activated NK cells are known to express PD-1, demonstrating that the observed pattern of PD-1 expression is consistent with the reduced ability of scRNA-seq to capture low to moderate expressed genes.

**Supplemental Figure 4.**
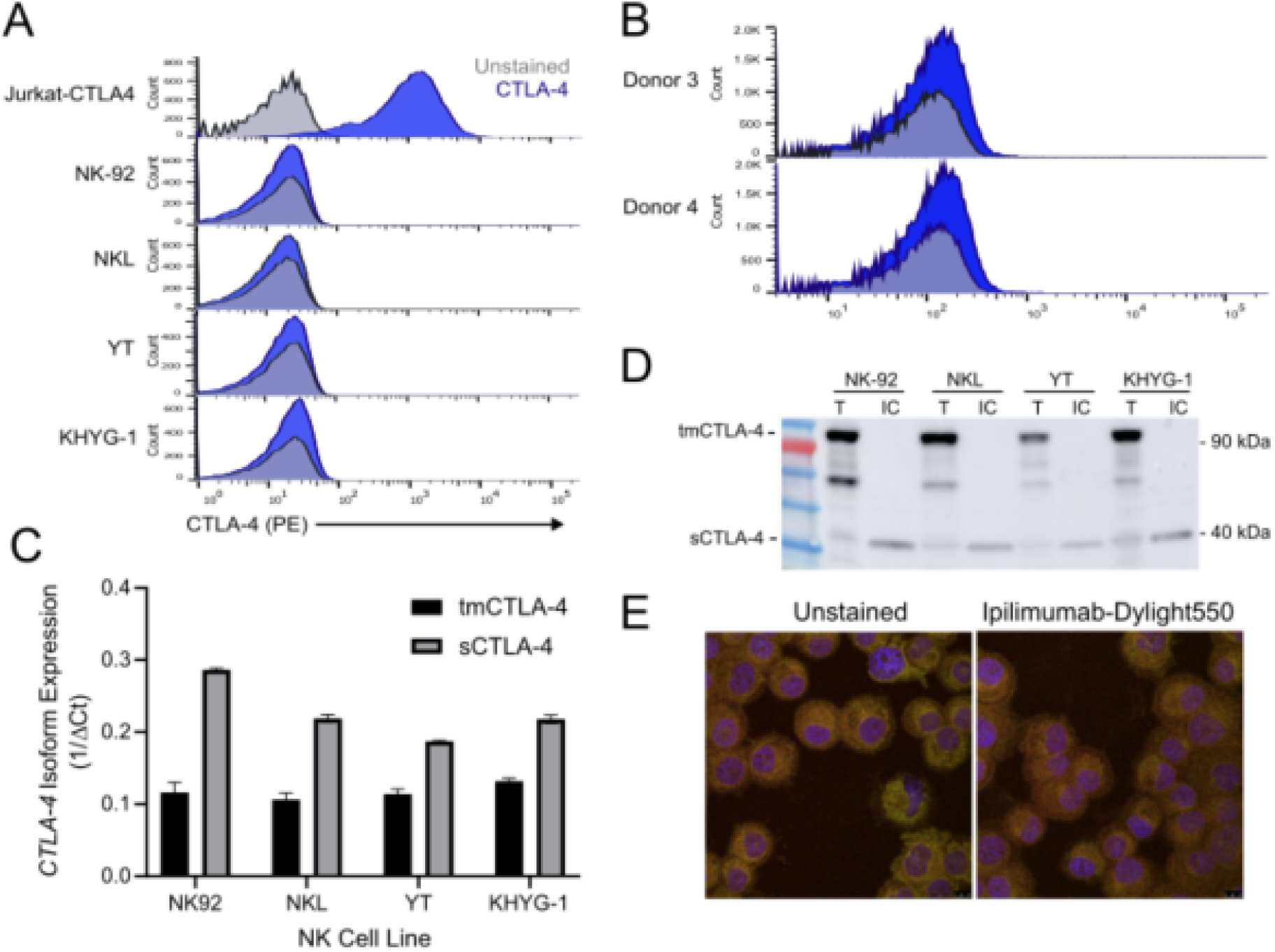
Human NK cell express CTLA-4. A. Flow cytometry for surface expression of CTLA-4 in positive control (Jurkat-CTLA4) and NK cell lines (NK-92, NKL, YT, KHYG-1). B. Flow cytometry for surface expression of CTLA-4 on CD56+ selected *ex vivo* unstimulated NK cells derived from healthy human donors C. Quantitative qRT-PCR analysis of transmembrane (tmCTLA-4) and soluble (sCTLA-4) isoforms in human NK cell lines. D. Western blot of total protein (T) and intracellular (IC) protein isolated from human NK cell lines NK-92, NKL, YT and KHYG-1 using cell surface protein biotinylation for exclusion of surface proteins demonstrating surface expression of CTLA-4 dimers and intracellular expression of CTLA-4 monomers. E. Immunofluorescent images of PANC-1 cells stained with Dylight550-labelled ipilimumab. Blue staining indicates DAPI. Shown are representative images of a single field of view taken via confocal microscopy (magnification, 63X).

**Supplemental Figure 5.**
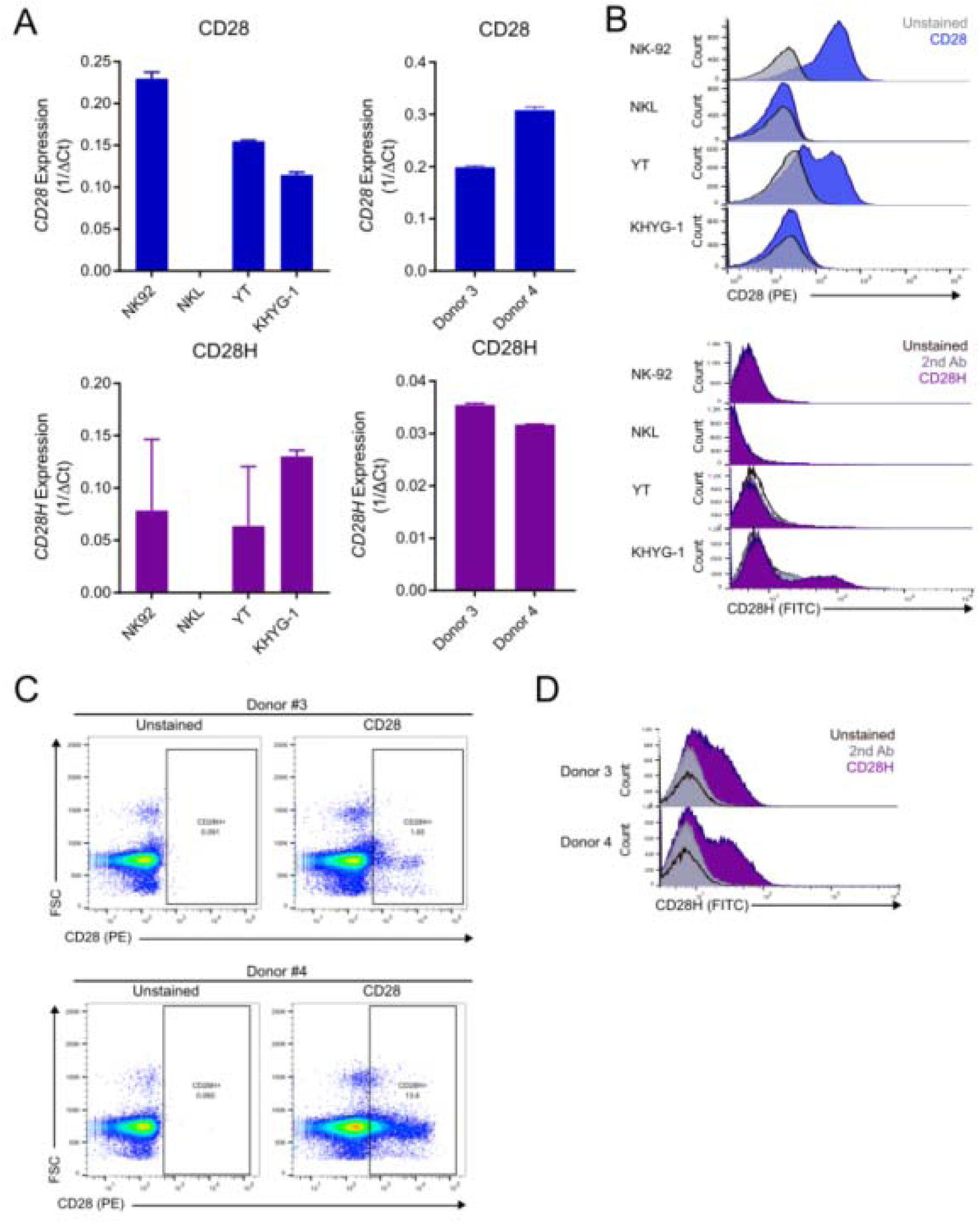
CD28 and CD28H expression on human NK cells. **A.** qRT-PCR assessment of CD28 and CD28H expression in human NK cell lines and primary donor NK cells. **B.** Flow cytometry assessment of CD28 and CD28H surface expression by human NK cell lines **C.** Flow cytometry assessment of CD28 surface expression by primary donor NK cells. **D.** Flow cytometry assessment of CD28H surface expression by primary donor NK cells.

**Supplemental table 1:** Gene set statistics for all 21 CoGAPS patterns

**Supplemental table 2:** Differentially expressed genes across pseudotime in NK cells collected from tumors treated with anti-CTLA-4

**Supplemental table 3:** Correlation values and p-values for cibesort cell type estimation and CTLA-4 expression in tumors from TCGA.

## Notes

**Disclosure of Potential Conflicts of Interest:** Elana J Fertig is on the Scientific Advisory Board of Viosera Therapeutics.

### Competing Interest Statement

Elana J Fertig is on the Scientific Advisory Board of Viosera Therapeutics.

